# Super-Resolved FRET and Co-Tracking in pMINFLUX

**DOI:** 10.1101/2023.03.24.534096

**Authors:** Fiona Cole, Jonas Zähringer, Johann Bohlen, Tim Schröder, Florian Steiner, Fernando D. Stefani, Philip Tinnefeld

**Author notes:** These authors contributed equally (Fiona Cole, Jonas Zähringer).

## Abstract

Single-molecule FRET (smFRET) is widely used to investigate dynamic (bio) molecular interactions taking place over distances of up to 10 nm. With the advent of recent super-resolution methods such as MINFLUX, MINSTED or RASTMIN, the spatiotemporal resolution of these techniques advanced towards the smFRET regime. While these methods do not suffer from the spatial restriction of FRET, they only visualize one emitter at a time, thus rendering fast dynamics of interactions out of reach. Here, we describe two approaches to overcome this limitation in pMINFLUX using its intrinsic fluorescence lifetime information. First, we combined pMINFLUX with smFRET. This enabled us to track a FRET donor fluorophore and simultaneously colocalize its FRET acceptor with nanometer precision. To extend co-localized tracking beyond the FRET range, we developed pMINFLUX lifetime multiplexing, a method to simultaneously track two fluorophores with similar spectral properties but distinct fluorescence lifetimes. We demonstrated its application on both static and dynamic DNA origami systems with a precision better than 2 nm. Within the FRET range, pMINFLUX lifetime multiplexing additionally uses a novel combined phasor-microtime-gating approach. This paves the way for nanometer precise co-localized tracking for inter-dye distances between 4 nm and 100 nm, closing the resolution gap between smFRET and co-tracking.

**Graphical Abstract:** 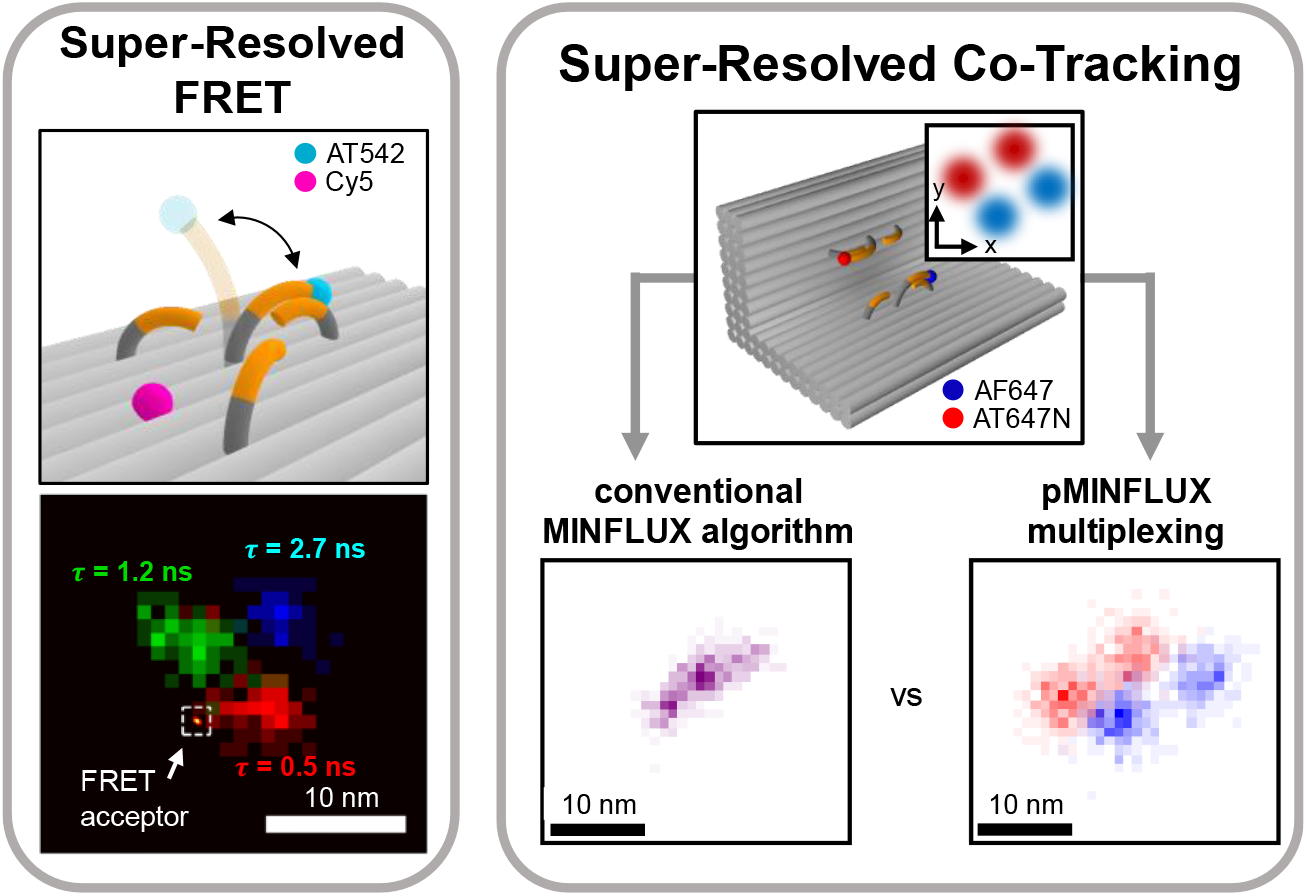

## Introduction

Molecular interactions and changes of conformational states are beautifully revealed with single-molecule FRET (smFRET).^1^ With its high sensitivity for small distance changes, smFRET has provided unique insight into the molecular mechanisms of life, including DNA replication^2,3^, transcription^2,4^, translation^5,6^ and repair^3^, protein folding^7,8^ and various enzymatic functions^9,10^. However, its working range limited to distances smaller than 10 nm leaves many other relevant biological dynamics occurring at larger distances, such as protein-protein interactions in dimerization, oligomerization and in multicomponent molecular machineries, out of reach for smFRET. Here, the co-tracking of multiple molecules of interest with nanometer precision may offer an alternative. However, experimental limitations such as chromatic aberrations in multicolor experiments and photon-inefficient single-molecule localizations by camera-based systems have so far restricted widespread applications. Combined singlemolecule tracking and FRET visualizations have therefore been rare (see e.g. refs ^11– 14^).

In recent years, new conceptual and technological advances including MINFLUX^15–17^ and later MINSTED^18^ and RASTMIN^19^ have demonstrated the routine localization of single-molecules with nanometer precision, thus reaching the working range of FRET. In MINFLUX, emitters are localized by sequentially interrogating their position with spatially displaced, vortex shaped excitation beams. By comparing the number of photons emitted upon excitation with the different beams with their known beam profiles, the absolute position of emitters is estimated. Furthermore, MINFLUX can reach sub-millisecond temporal resolution ideal for tracking applications with strikingly optimized photon budgets.^20,21^ However, while multiplexing with MINFLUX was achieved in imaging by spectral splitting^17^ and Exchange DNA-PAINT^22^, fast simultaneous tracking below the diffraction limit with nanometer precision remained restricted to single emitters. As soon as two or more fluorescent emitters are present, neither of them could be localized correctly, making multiplexed tracking experiments so far unfeasible with these techniques.

Here, using pulsed interleaved MINFLUX (pMINFLUX)^16,23^ in a series of experiments on DNA origami model nanostructures, we demonstrate multiplexed single-molecule tracking spanning from the FRET range to the range of super-resolved tracking. pMINFLUX provides fast localization rates by performing the excitation sequence at the repetition rate of the laser, typically in MHz and direct access to the fluorescence lifetime.^16^ The fluorescence lifetime information both enables additional FRET efficiency determination within the FRET range and serves as a distinguishable characteristic to separate the photons from different molecules as previously demonstrated in FLIM and STED microscopy.^24–26^

In a first experiment, we tracked a FRET donor labelled DNA sequence which transitions between the vertices of a triangular structure with 6 nm side length. Tracking with pMINFLUX provides the donor position and at the same time detects the proximity of a FRET acceptor by the shortened fluorescence lifetime of the donor. Calculation of the FRET efficiency enables the estimation of the inter-dye distance at the different donor positions. This, in turn, enables determining the position of the acceptor relative to the triangular structure. Next, we introduce a concept to localize two dye molecules with similar spectra but distinct fluorescence lifetimes in distances beyond the FRET range without photoswitching. We simultaneously localize an ATTO647N molecule and an Alexa Fluor 647 molecule separated by 15 nm with a precision only slightly reduced compared to individual localizations. We then continue to use this concept to simultaneously track the position of two dye molecules that independently jump between different sites on a DNA origami, demonstrating its applicability in superresolved co-tracking beyond the FRET range. With a combined phasor-microtime gating approach, we extend the concept to dye molecules positioned at distances within the FRET range and thus close the resolution gap between single-molecule FRET and co-tracking. Altogether, these experiments exemplify how MINFLUX in combination with the fluorescence lifetime information can provide new insights into molecular interactions and dynamics occurring in the FRET range and beyond.

## Results

### Super-Resolved FRET in pMINFLUX

We used a DNA origami molecular balance^27^ as a platform to carry out simultaneous p-MINFLUX and FRET tracking experiments (Fig. 1A). The molecular balance features a 19 nucleotides (nt) long single-stranded DNA pointer labelled at the tip with an ATTO542 molecule (FRET donor). The position of the donor was tracked via pMINFLUX using green excitation (Supplementary Fig. 1). The pointer can transiently hybridize to three single-stranded protrusions placed in a nearly equilateral triangle of ∼6 nm side length, via an 8 nt complementary sequence. An additional Cy5 molecule (FRET acceptor) was placed at a fixed position in proximity to the protrusions on the DNA origami structure, (details in Supplementary Tables 2-4).

**Figure 1.**
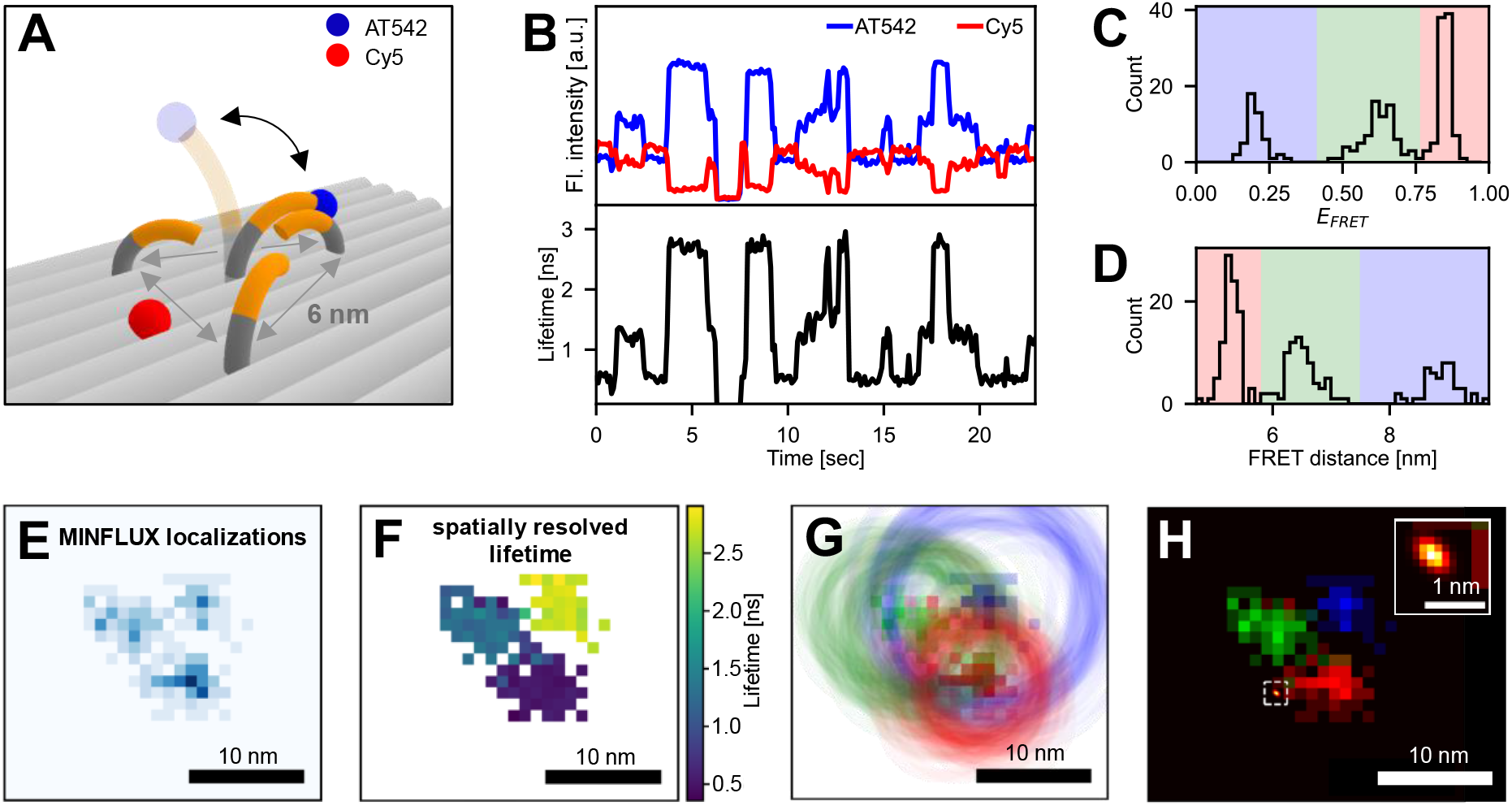
Super-resolved FRET in pMINFLUX. (A) Schematic of the dynamic DNA origami with three protruding strands in distances of 6 nm to each other to which an ATTO542 labelled DNA pointer transiently hybridizes. (B) Anticorrelated fluctuations in ATTO542 (blue) and Cy5 (red) fluorescence between three intensity levels which are correlated to fluctuations in the fluorescence lifetime of ATTO542, indicating transitions of the DNA pointer between three positions. (C, D) FRET efficiency and FRET distance distributions calculated from the fluorescence lifetimes of ATTO542, featuring three distinct populations highlighted in blue, green and red. (E) 2D histogram of the pMINFLUX localizations of the DNA pointer. (F) Spatially resolved fluorescence lifetimes of the ATTO542 dye on the DNA pointer. (G,H) Multilateration of the position of the Cy5 dye. By combining each MINFLUX localization of the DNA pointer (square) with a FRET distance (circle) (G), a multiplicative probability density map of the position of the Cy5 molecule is constructed (H). The inset shows a zoom-in around the area of the multilaterated Cy5 position highlighted by the dashed white box. The corresponding maximum in the probability density map has a full-width at half-maximum of only FWHM = 0.17 nm.

To monitor FRET, the detection was spectrally split in two channels for donor and acceptor emission. The fluorescence transients recorded for ATTO542 and Cy5 show anticorrelated fluctuations between three intensity levels which are correlated with fluctuations of the fluorescence lifetime of ATTO542 (Fig. 1B), as expected for the DNA pointer transitioning between the three positions. Calculation of the FRET efficiency from the fluorescence lifetime of ATTO542 (Fig. 1C) reveals that the donor-acceptor distances for the three pointer positions are 5.3 nm, 6.7 nm and 9.5 nm (Fig. 1D, R_0_ = 7.1 nm, Supplementary Note 2).

Localization analysis of the same pMINFLUX transient shown in Fig. 1B delivers the binding positions of the donor with ∼1-2 nm precision at 100 ms temporal resolution. The two-dimensional localization histogram shown in Fig. 1E confirms that the DNA pointer visited the three positions separated ∼6 nm from each other, in agreement with the DNA origami design.

For synergistically combining MINFLUX with FRET, each MINFLUX localization was assigned to its corresponding fluorescence lifetime. The resulting super-resolved fluorescence lifetime image in Fig. 1F shows that the ATTO542 molecule decays with different lifetimes depending on the pointer position, confirming that both FRET and MINFLUX data describe the DNA pointer dynamics well.

Next, both methods were combined to maximize their information output. From each pMINFLUX localization, both the position of ATTO542 and its separation distance to Cy5 were determined (the latter using the fluorescence lifetime information). Then, a circle centered at the ATTO542 position with a radius of the assigned FRET distance was defined. Doing this repeatedly as the DNA pointer explores the three positions delivered three sets of circles (Fig. 1G, Supplementary Video 1). From these circles, a multiplicative probability density map for the location of Cy5 was created (Fig. 1H, Supplementary Fig. 2). This density map featured a single, very narrow peak with a full-width at half-maximum of FWHM = 0.17 nm, determining the position of the Cy5 molecule in close proximity to two of the three DNA pointer locations by multilateration.^28–31^

As such, we demonstrated the combination of pMINFLUX localizations with their intrinsic FRET information. Besides applications in multilateration approaches, this combination can generally be used to track the absolute position of a molecule with pMINFLUX while simultaneously tracking the position of a second molecule relative to the first molecule using FRET. Due to the simplicity of its experimental implementation on pMINFLUX setups, we believe that this combination has the potential to become a powerful tool when studying biological interactions. However, due to the use of FRET as an information source, the approach probes interactions occurring in distances of below ∼12 nm. If molecular interactions occur with the dyes being further apart, structural information is lost. Thus, other methods that exploit the optical distinguishability of different emitters either spectrally or in the fluorescence lifetime domain are required for MINFLUX.

### Fluorescence Lifetime-Multiplexing for Co-Tracking in pMINFLUX

To this end, we developed a fluorescence lifetime based multiplexing of pMINFLUX to localize more than one emitter simultaneously without photoswitching. pMINFLUX lifetime multiplexing is based on obtaining the fluorescence intensities necessary for the position estimation from fits to the fluorescence decays corresponding to each one of the four excitation beams rather than from the conventional counting of the absolute number of photons emitted upon excitation with the four different beams (Supplementary Section 3). A first test of the suitability of this approach was done by comparing pMINFLUX localizations of single AlexaFluor647 (AF647) molecules obtained from intensity counts and from monoexponential fits. Localizations with both approaches exhibit negligible differences in AF647 position and localization precision (Supplementary Fig. 3).

Next, we set out to localize two emitters simultaneously using lifetime multiplexing. For these experiments, we designed a static DNA origami with two fluorophores, an AF647 and an ATTO647N molecule, placed in fixed positions with a nominal separation distance of 14.6 nm (Fig. 2A). AF647 and ATTO647N have similar spectral properties (see Supplementary Fig. 4), but distinct fluorescence lifetimes of 1.1 ns and 4.3 ns, respectively. Figure 2B shows a fluorescence intensity transient with two photobleaching steps recorded for a single DNA origami structure in a pMINFLUX measurement. The fluorescence lifetime decay before the first bleaching step exhibits a biexponential behavior, indicating the presence of both dyes (time window I in Fig. 2B). After the first photobleaching event, the decay shows a monoexponential profile with a fluorescence lifetime of 4.3 ns corresponding to ATTO647N (time window II in Fig. 2B).

**Figure 2.**
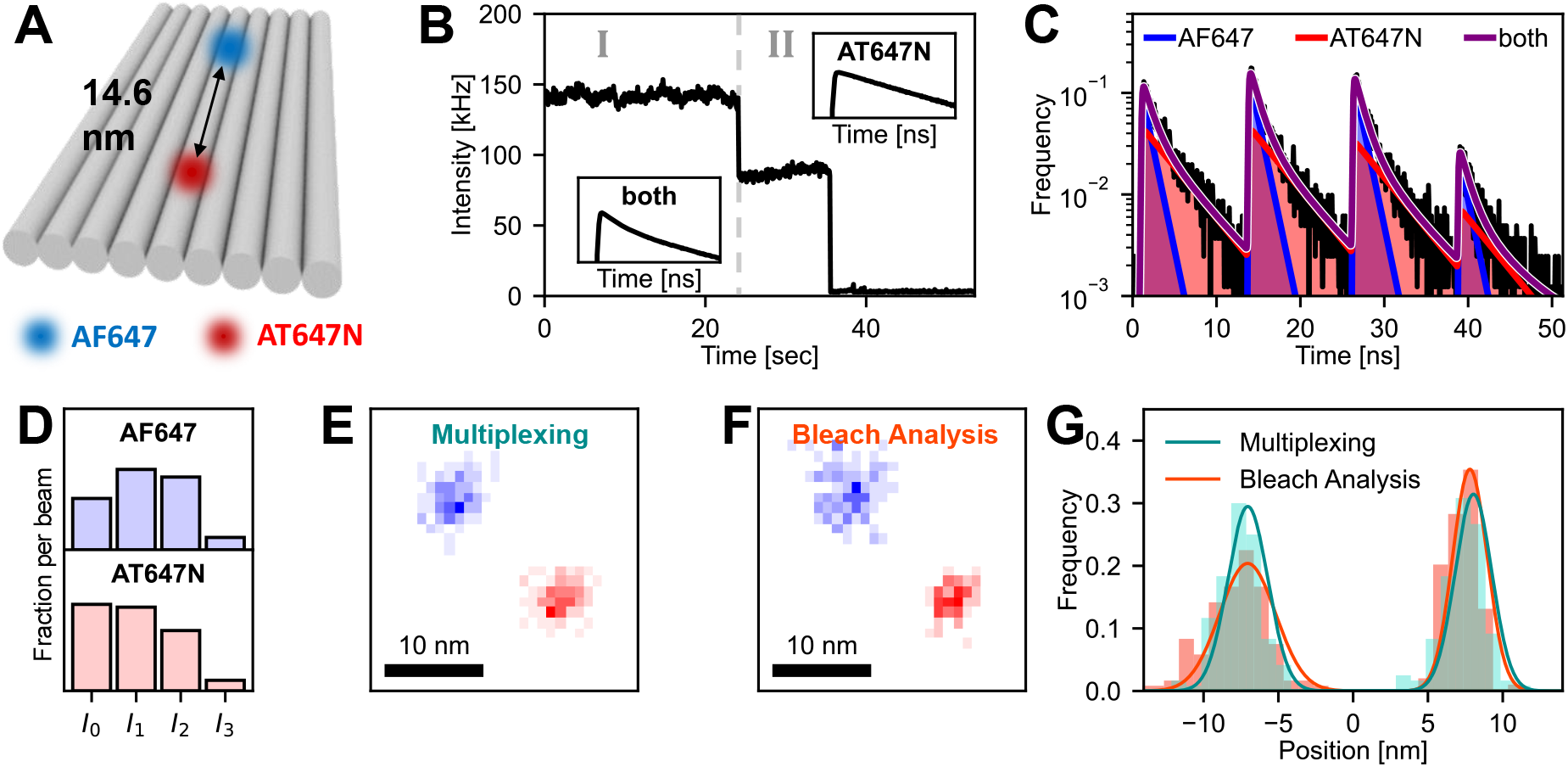
pMINFLUX lifetime multiplexing principle and accuracy. (A) Schematic of a static DNA origami with AF647 and ATTO647N placed in a fixed distance of ∼14.6 nm from each other. (B) Fluorescence intensity transient recorded for a single DNA origami structure shown in panel A during a pMINFLUX measurement. Insets show the fluorescence lifetime decays before and after photobleaching of AF647 (dashed gray line). (C) Fluorescence microtime decays for time window I in panel B. Biexponential fitting (purple) reveals the fluorescence decay profiles of AF647 (blue) and ATTO647N (red) separately. (D) Relative fluorescence intensities recorded upon excitation from the four pulsed-interleaved beams for both dyes. The intensity values were extracted from the biexponential fit model shown in Panel C. (E) 2D histogram of the lifetime multiplexed pMINFLUX localizations recorded while both dyes simultaneously were in their fluorescent state. Localizations of AF647 and ATTO647N are shown in blue and red, respectively. (F) 2D histogram of pMINFLUX localizations of the same trace obtained by bleach analysis (see Supplementary Fig. 6). (G) Line profiles of the localizations shown in Panel E,F projected along the axis of both localizations. The two maxima found with both approaches correspond to the localizations of AF647 (left) and ATTO647N (right).

Measurement time window I where both fluorophores were in their emissive state (Supplementary Fig. 5) was analyzed using a biexponential fit. From the fit, the fluorescence decays of AF647 and ATTO647N were extracted (Fig. 2C), to obtain their fluorescence intensities upon excitation with each one of the four beams (blue and red overlays in Fig. 2C,D). The resulting two-dimensional localization histogram features two distinct populations describing the positions of AF647 and ATTO647N, with a separation distance of 15.0 nm in agreement with the DNA origami design (Fig. 2E).

We tested the accuracy of the lifetime multiplexing by comparing its results to localizations obtained using the so-called bleach analysis. Analogous to similar approaches used in wide-field super-resolution imaging,^32–34^ we performed measurements until both molecules photobleached. Then, we first localized the lasting molecule (ATTO647N) after the first photobleaching event, as shown in Fig. 2B, using the data in time window II. The position of the other molecule (AF647) was estimated by subtracting the average fluorescence intensity of ATTO647N from the fluorescence of both dyes in time window I (Fig. 2F, Supplementary Fig. 6). The resulting positions of both dyes match the positions obtained by lifetime multiplexing (Fig. 2G). In contrast to lifetime multiplexing, the bleach analysis can only be applied to static systems.

Next, we evaluated the localization precision achievable in lifetime multiplexed pMINFLUX. Using the same measurement shown in Figure 2, we varied the number of photons used for each localization. Figures 3A-F depict two-dimensional histograms of lifetime multiplexed localizations of AF647 and ATTO647N performed with photon counts between 100 and 4000. The localization precision of both molecules follows the expected inverse dependency with √*N*, and with moderate photon counts of 2000 photons per emitter, both emitters were simultaneously localized with precisions better than 3 nm.

**Figure 3.**
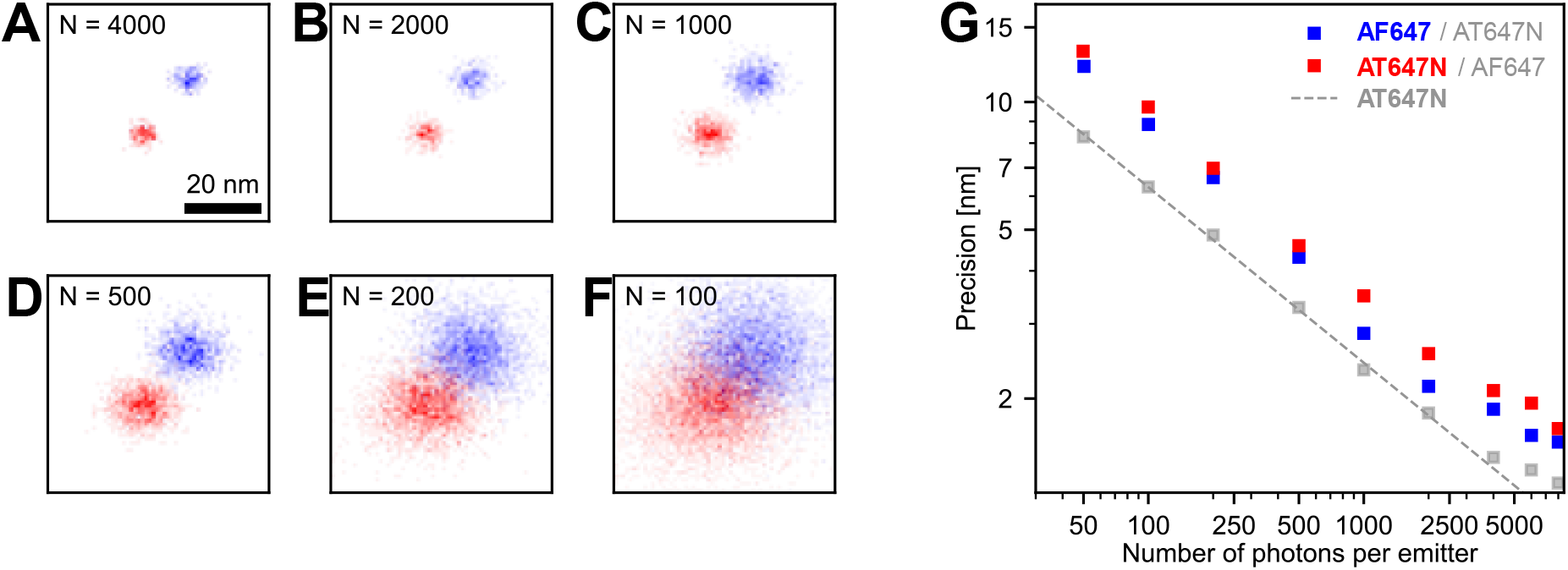
Evaluation of the localization precision in pMINFLUX lifetime multiplexing. (A-F) 2D histograms of the lifetime multiplexed pMINFLUX localizations of an AF647 (blue) and an ATTO647N (red) dye in a fixed distance of 14.6 nm for different number of photons per emitter, N, used to estimate their positions. (G) localization precision as a function of collected photons per emitter for both AF647 (blue) and ATTO647N (red) localized simultaneously in pMINFLUX lifetime multiplexing. The localization precision of a single ATTO647N dye at the same position as in the multiplexing localized after photobleaching of AF647 is shown in gray.

Interestingly, the multiplexed simultaneous localizations have precisions about 40% worse compared to the localization precision attained for the individual ATTO647N molecule (time window II in Fig. 2B), (Fig. 3G). These lower precisions can be attributed to uncertainties of photon assignment. To investigate this, we performed numerical simulations (Supplementary Section 8) varying the relative intensity and lifetime contrast of the two target fluorophores. We found that the brightness ratio of the emitters does not have a systematic influence on the attainable localization precision. By contrast, the localization precision reduces with lifetime contrast (Supplementary Fig. 7), which explains the reduction in localization precision observed in the experiments for the pair AF647-ATTO647N. Using the simulation framework, other suitable dye pairs for lifetime multiplexed pMINFLUX can be identified. As example, we identified and experimentally demonstrated the suitability of ATTO542 and Alexa Fluor 555 to expand the range of multiplexed pMINFLUX to the green range (Supplementary Fig. 8).

In contrast to conventional MINFLUX nanoscopy, lifetime multiplexed pMINFLUX offers the possibility to track multiple emitters simultaneously on the nanoscale. To demonstrate its potential in tracking applications, we designed a dynamic DNA origami, similar to the FRET pointer system shown Figure 1A, but in this case the structure features two DNA pointers, one labelled with AF647 and the other with ATTO647N (Fig. 4A). Each DNA pointer can transiently hybridize to two single-stranded protrusions on the DNA origami distanced ∼12 nm from each other. By choosing different lengths for the sequences complementary to the DNA pointer on the protrusions, differing kinetics for each DNA pointer system were achieved.^27^

**Figure 4.**
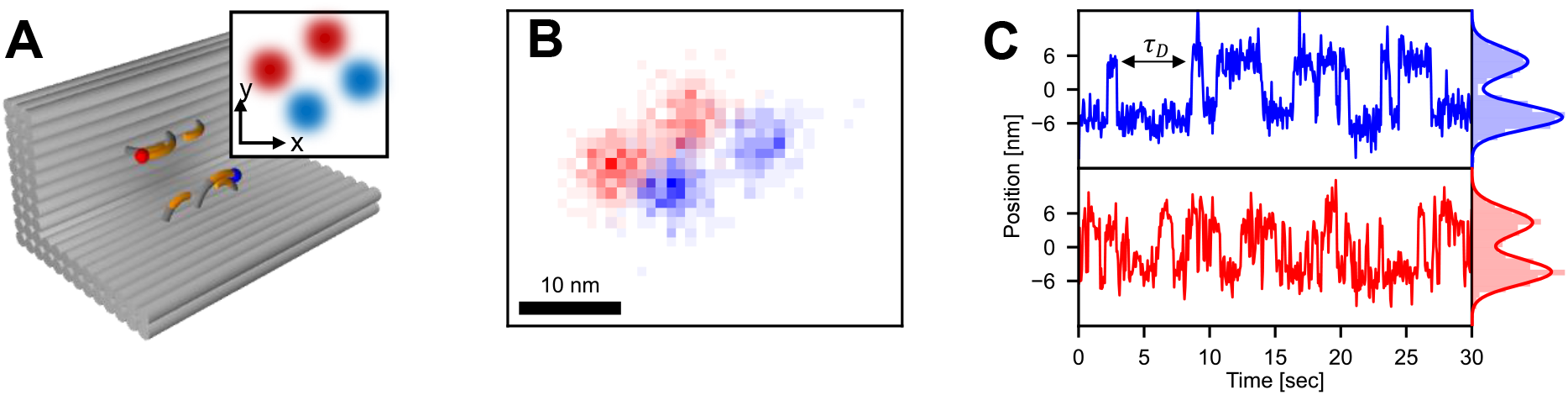
Simultaneous tracking using lifetime multiplexed pMINFLUX. (A) Schematic of the dynamic DNA origami with both an AF647 labelled DNA pointer (blue) and an ATTO647N labelled DNA pointer (red) which can each transiently hybridize to two protruding strands distanced ∼12 nm from each other. The pointer strands have complementary sequences of 8 nt and 7 nt to the protruding strands for the AF647 and the ATTO647N pointer, respectively. The inset shows the xy-projection of the protruding strands to which AF647 (blue) and ATTO647N (red) can bind. (B) 2D histogram of the lifetime multiplexed pMINFLUX localizations of AF647 and ATTO647N, featuring each two distinct positions. (C) Localization trajectory of the AF647 (blue) and ATTO647N DNA pointer (red), revealing uncorrelated fluctuations between two positions with different kinetics for both DNA pointers. The double-headed arrow indicates one dwell time *τD* the AF647 pointer spends bound to one protruding strand.

Lifetime multiplexed pMINFLUX enabled the simultaneous tracking of both fluorophores as they jump between the two binding positions (Fig. 4B,C, Supplementary Video 2). The two-dimensional localization histogram (Fig. 4B) shows that each fluorophore explores two positions in agreement with the designed geometrical arrangement. Kinetics of the transitions between the different protrusions were extracted from spatial trajectories of the DNA pointers, separately for each dye. The AF647 DNA pointer system with complementary sequence of 8 nt with the docking site shows a mean dwell time of *τ*_*D*_ = 1.5 s at each protrusion, whereas the ATTO647N system with a shorter complementary sequence of 7 nt exhibited a dwell time of *τ*_*D*_ = 0.5 s. Due to the differing kinetics, the measurement data was re-analyzed with different temporal resolutions individually for both DNA pointers to achieve the best trade-off between temporal resolution and localization precision, separately for both systems (Supplementary Fig. 9). The flexibility of optimizing spatio-temporal resolution in post-processing is another advantage of pMINFLUX lifetime multiplexing.

### pMINFLUX Lifetime Multiplexing within the FRET Range

At distances compatible with FRET, fluorophores with similar spectra such as AF647 and ATTO647N interact and cannot be considered independent from each other. Due to the mutual overlap of excitation and emission spectra of both fluorophores, FRET occurs both from AF647 to ATTO647N and vice versa. As a consequence, a detected photon cannot accurately be assigned to the excitation of a specific fluorophore, as needed for MINFLUX localizations (Fig. 5A). Instead, as the distance shortens and FRET becomes stronger, the fluorescence decay shows an increasingly monoexponential profile (see Fig. 5B, Supplementary Fig. 10). Naturally, this affects the accuracy of the lifetime multiplexing. Monte Carlo simulations for AF647 and ATTO647N at different distances to each other show deviations from the estimated distances to the ground truth at inter-dye distances smaller than ∼10 nm (see Fig. 5C).

**Figure 5.**
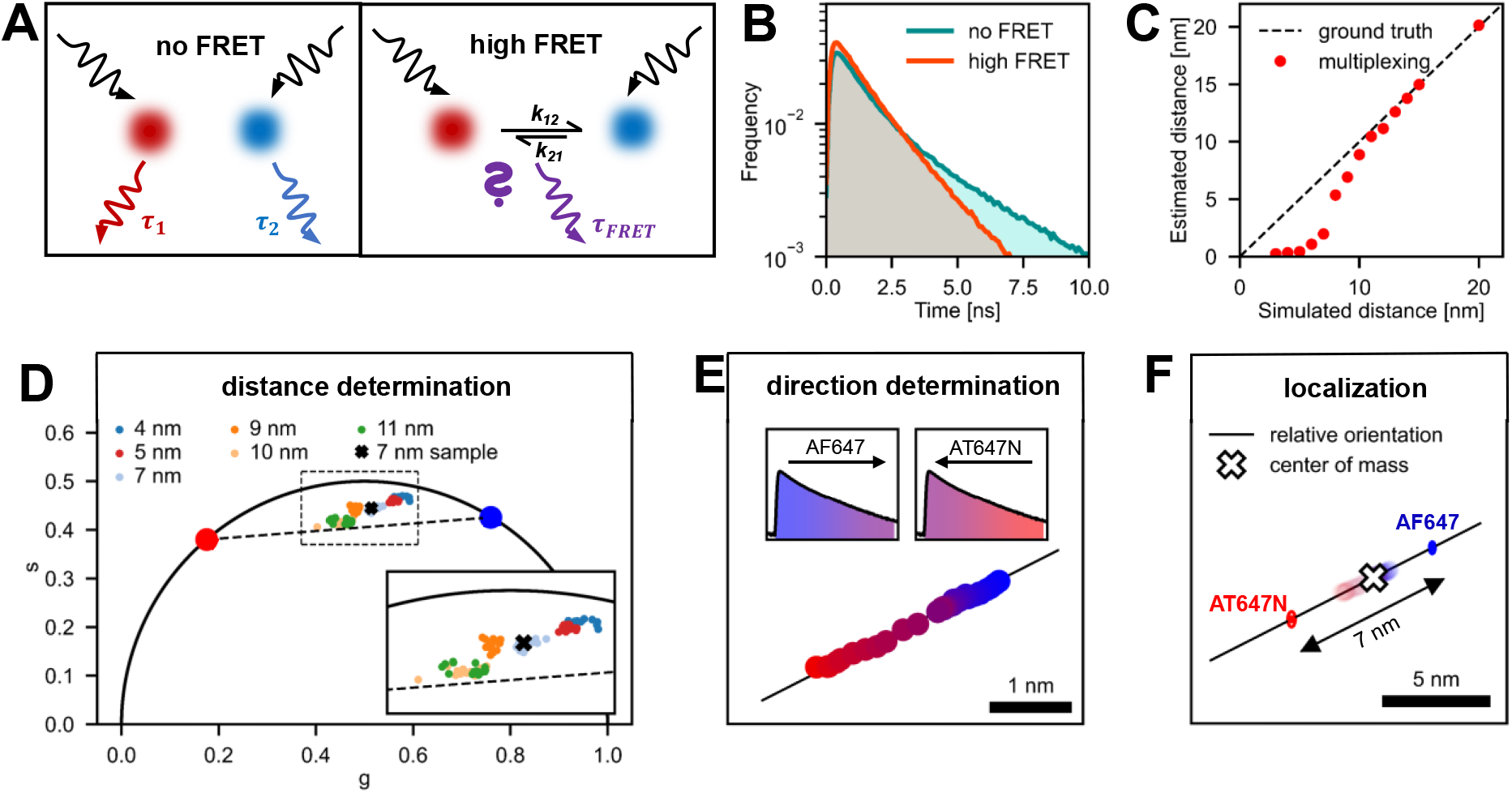
pMINFLUX lifetime multiplexing within the FRET range. (A) At large distances when no FRET occurs, the fluorescence of two emitters can be considered to be independent of each other. Photon absorption and emission take place at the same emitter. At small distances, the fluorescence of two emitters with similar spectral properties becomes coupled. Due to FRET in both directions (k12, k21), photons are not necessarily emitted from the emitter that absorbed the photon. (B) Experimental fluorescence lifetime decays measured for the combination of AF647 and ATTO647N at a distance of 14 nm (no FRET) and a distance of 4 nm (high FRET). (C) Accuracy of pMINFLUX lifetime multiplexing in the FRET range according to Monte-Carlo simulations. The dashed black line has a slope of one and indicates perfect accuracy. (D) Phasor plot of DNA origami structures with AF647 and ATTO647N placed at different distances. The red and blue dots correspond to the fluorescence of pure ATTO647N and AF647, respectively. Datapoints on the dashed black line between them indicate structures containing both dyes in distances without interactions. The inset shows a zoom-in of the area highlighted by the dashed box. The black cross corresponds to the measurement used to demonstrate the phasor/ microtime gating based localization approach in (E,F). (E) pMINFLUX microtime gating in the FRET range. By analyzing only a subset of photons selected in microtime gate windows, the direction defined by the positions of AF647 and AT647N can be determined. Increasing the size of the microtime gate from early (blue) to late detected photons (purple) after pulsed laser excitation (left inset) and from late (red) to early detected photons (right inset) yields a line of localizations along the line defined by the positions of the two target molecules (black line). The corresponding localizations are shown with a color gradient from blue to purple for microtime gates of increasing size for the early photons and from red to purple for microtime gates of increasing size for the late photons. The color code corresponds to the color gradient used to visualize the gradual expansion of the microtime gates in the insets. (F) pMINFLUX position estimation in the FRET range. Combining the distance information from the phasor plot (black arrow) with the found distance from the microtime gating (black line) and the center of mass localization of the coupled system (white cross) yields an estimation on the absolute position of AF647 and ATTO647N (blue and red ellipses).

To achieve accurate simultaneous localizations with pMINFLUX in the FRET range, we complemented the lifetime multiplexing with phasor analysis^35^ and microtime-gated detection^27,36^. The idea is that, although the positions of the fluorophores cannot be determined directly, they can be deduced from their separation distance, the direction of the connecting vector and the center of mass of the two positions.

The separation distance can be obtained using the phasor approach.^35^ Under this framework, emitting species with a pure monoexponential decay have a phasor lying along a so-called universal circle. As the coordinates of the phasor plot are additive, the phasor of systems of two fluorophores in which no FRET occurs lie on the line joining the individual phasors, in our case the ones of the AF647 and the ATTO647N phasor (Fig. 5D). In such systems, lifetime multiplexed pMINFLUX works well as described above. By contrast, if FRET occurs, the resulting phasor deviates from this line. In our case of mutual FRET, as the separation shortens, the coupling between the two fluorophores increases and eventually they display an increasingly monoexponential decay similar to a single emitter. In the phasor analysis this shows as a deviation from the AF647-ATTO647N lines towards the universal circle (Fig. 5D, Supplementary Section 11). Using a calibration with DNA origami structures containing AF647 and ATTO647N at different fixed distances, this deviation can be calibrated to estimate the separation distance between AF647 and ATTO647N in the FRET range (see Fig. 5D, Supplementary Fig. 11).

The direction defined by the actual positions of AF647 and ATTO647N is then determined by combining the standard pMINFLUX localization algorithm with microtime gated detection^27,36^ (Fig. 5E). Due to the different fluorescence lifetimes of AF647 and ATTO647N, photons arriving shortly after each excitation pulse are predominantly due to AF647 emission, whereas photons with late microtimes are mainly emitted from ATTO647N. Thus, performing pMINFLUX using microtime gates at the beginning and at the end of each excitation window leads to inaccurate localizations at intermediate positions between the two molecules. Gradually increasing the size of the microtime gates gives a sequence of localizations along the line defined by the positions of the two target molecules.

Finally, performing a localization using all detected photons (no microtime gating) delivers the center of mass of the coupled system of both fluorophores. Combining this center of mass with the separation distance between both fluorophores as determined by phasor analysis and the direction defined by their positions extracted from the microtime gating approach yields two absolute positions for AF647 and ATTO647N (Fig. 5F), extending the applicability of our pMINFLUX lifetime multiplexing approach to the FRET range.

We additionally validated the accuracy of the phasor/microtime-gating based localization approach in the FRET range by performing Monte-Carlo simulations for systems with AF647 and ATTO647N placed at different distances. At distances above ∼ 4 nm, the approach localizes both fluorophores accurately. At lower distances, high FRET values reduce the distinguishability of both fluorophores, resulting in a lower colocalization bound of about 4-5 nm (see Supplementary Fig. 10-12).

In analogy to the pMINFLUX lifetime multiplexing approach in the fluorescence lifetime domain, we propose a multiplexing approach based on differing spectral properties of emitters. By exploiting small shifts in the emission spectra of different emitters, their fluorescence responses are also separated without photoswitching, resulting in their simultaneous localization (see Supplementary Fig. 13). This alternative approach can also be applied to continuous-wave MINFLUX. With the emitter pair ATTO542-Cy3B which exhibits good photostability ideal for co-tracking, it yields localization precisions only two-fold worse compared to single emitters in simulations. While this is slightly worse than the decrease in localization precision by a factor of 1.4 experimentally observed in pMINFLUX multiplexing in the lifetime domain, it still suffices for photo-efficient tracking. In future, a redundancy achieved by combining both approaches can be used to robustly apply simultaneous multiplexing in more complex environments such as cells.

## Conclusion

In conclusion, we demonstrated how the fluorescence lifetime information intrinsic to pMINFLUX measurements is synergistically used to combine MINFLUX localizations with FRET. The combination allows tracking single emitters on the nanoscale while simultaneously scanning their immediate environment for the presence of acceptor molecules, the distance to which can also be determined. Next, we established an approach to track multiple emitters simultaneously in pMINFLUX using only one excitation color by separating them based on their fluorescence lifetimes. At distances above 10 nm, the position of two emitters is estimated with nanometer precision by implementing a biexponential fluorescence lifetime fit in the pMINFLUX localization algorithm. At lower distances, a combined phasor-microtime gating approach allows their simultaneous localization.

Both the combination of pMINFLUX and FRET as well as the pMINFLUX lifetime multiplexing approach are not limited to pMINFLUX. As all developed algorithms are based solely on the microtime information of the emitters, no further instrumental effort is required. As such, both approaches have great potential to become generalized techniques for the simultaneous super-resolved tracking of two or more emitters. Their implementation should in principle be possible in any stochastic super-resolution technique with fluorescence lifetime information, such as RASTMIN,^19^ confocal fluorescence-lifetime single-molecule localization microscopy^37^ and wide-field fluorescence lifetime imaging.^38^ Also, the principal findings are not limited to twodimensional imaging as shown here but should be extendable to 3D super-resolution microscopy.

We additionally developed a multiplexing approach in the spectral domain which also allows the simultaneous localization of two emitters for setups operating with continuous-wave lasers. Here, the same spatial limitations as for the pMINFLUX lifetime multiplexing approach hold: FRET sets a lower limit for the resolution at ∼4 nm below which photons originating from emitters become indistinguishable. An upper limit is given by the limited field of view of MINFLUX. Depending on the spatial arrangement of the excitation beams, molecules in distances of up to 100 nm can be tracked simultaneously. Similar to all MINFLUX experiments, this field of view could be further extended by the use of single-photon avalanche detector arrays,^39^ yielding the simultaneous nanometer precise localizations of multiple emitters over a large range of distances. Overall, we envision that further developing these approaches will pave the way for nanometer precise multi-color tracking experiments in living cells. The approaches could directly visualize dynamic process such as the stepping mechanism of kinesin motor proteins^20,40^ or diffusion processes through nuclear pore complexes on the nanoscale,^41^ giving more direct insights into dynamic processes in interplay with their environment in cell biology.

## Resource availability

### Lead contact

Philip Tinnefeld: Department of Chemistry and Center for NanoScience, LudwigMaximilians-Universität München, Butenandtstr. 5-13, 81377 München, Germany.

## Experimental methods and analysis

### Preparation of DNA origami structures

DNA origami structures were designed using the open-source software caDNAno2^42^ and assembled and purified using published protocols.^43^ Positions and distances of dyes in DNA origami structures were estimated assuming a distance of 0.34 nm between the nucleotides along the DNA double helix and 2.7 nm between the centers of adjacent helices.^44,45^ For the exact sequences of all unmodified and modified DNA staple strands used to fold the DNA origami structures see Supplementary Tables 2-7. DNA staple strands were purchased from Eurofins Genomics GmbH (Germany) and Integrated DNA Technologies (USA). p8064 scaffold used for the dynamic pointer origamis and p7049 used for the static two-color origamis (both derived from M13mp18 bacteriophages) were produced in house.

For DNA origami folding, 10 nM scaffold in 1xTAE (40 mM Tris, 20 mM acetic acid, 1 mM EDTA; pH 8) containing 12.5 mM/ 20 mM MgCl_2_ (static/ dynamic origami) was mixed with a 10-fold excess of all unmodified and a 30-fold excess of all modified oligonucleotides, respectively. The mixture was heated to 65 °C and kept at this temperature for 15 min before being cooled down to 25 °C either with a temperature gradient of -1 °C min^-1^ (static origami) or with a non-linear thermal annealing ramp over 16 h^46^ (dynamic origami). Folded DNA origami were purified from excessive staple strands by gel electrophoresis. All gels were ran using a 1.5% agarose gel, 1xTAE containing 12.5 mM MgCl_2_ for 2 hours at 6 V/cm. The target band containing DNA origami was cut from the gel and DNA origami solution extracted from the band via squeezing. Samples were stored at -20 °C until further use.

### Surface sample preparation for pMINFLUX measurements

As sample chambers, flow chambers consisting of a glass coverslip glued onto an objective slide with double-sided scotch tape were used. Prior to chamber assembly, coverslips were cleaned by incubation with 1% Hellmanex for 20 min followed by two 15 min washing steps with MilliQ water. After surface passivation by incubation with BSA-Biotin (0.5 mg/mL, Sigma Aldrich, USA) for 10 min, the surface was washed with 200 μL 1×PBS buffer (137 mM NaCl, 2.7 mM KCl, 10 mM Na_2_HPO_4_, 1.8 mM KH_2_PO_4_; pH 8). 150 μL neutrAvidin (0.25 mg/mL, Thermo Fisher, USA) was incubated for 10 min and then washed with 200 μL 1×PBS buffer. DNA origami solution was diluted in 1×TE buffer (10 mM Tris, 1 mM EDTA) containing 750 mM NaCl to a concentration of ∼100 pM and then immobilized on the biotin-neutrAvidin surface via biotin-neutrAvidin interactions. For this, 100 μL of the DNA origami sample solution was added and incubated for 5 min. Residual unbound DNA origami was removed by washing the chambers with 150 μL 1xTE buffer containing 750 mM NaCl. Next, gold nanorods with a longitudinal LSPR peak at 900 nm (fabricated following established protocols)^47^ were immobilized on the surface as fiducial markers for drift correction. Chambers were incubated with a diluted gold nanorod solution in 1xTAE containing 12.5 mM MgCl_2_ for 2 min and flushed with 150 μL 1xTAE (12.5 mM MgCl_2_). Directly prior to MINFLUX measurements, an oxidizing and reducing buffer system (1xTAE, 12.5 mM MgCl_2_, 2 mM Trolox/ Troloxquinone)^48^ was added in combination with an oxygen scavenging system (12 mM protocatechuic acid, 56 μM protocatechuate 3,4-dioxygenase from pseudomonas sp., 1% glycerol, 1 mM KCl, 2 mM Tris, 20 μM EDTA) to suppress blinking and photobleaching. After photostabilization chambers were sealed with picodent twinsil and measured. **pMINFLUX setup**

A description of the pMINFLUX setup is given in the first pMINFLUX implementation.^16^ For detailed information see Supplementary Section 1.

### Data analysis

Data processing and analysis of the MINFLUX experiments was realized using custom-written Python scripts. A description of the used algorithms is given in the Supplementary Information. All Python scripts used for data analysis are available from the authors upon request.

## Acknowledgments

P.T. gratefully acknowledges financial support from the Deutsche Forschungsgemeinschaft (DFG, German Research Foundation) – Project-ID 201269156 – SFB 1032 (A13), Ti329/15-1 (project number 470075523), from the Federal Ministry of Education and Research (BMBF) and the Free State of Bavaria under the Excellence Strategy of the Federal Government and the Länder through the ONE MUNICH Project Munich Multiscale Biofabrication and by G ‘ S g - X 89 -390776260. We thank Luna, Xaverl and Charly for their support with the measurements and data analysis and Angelika, Hildegard, Frau Steger and Herr Ehrl for lab upkeep.

## Author contributions

F.C., J.Z., T.S., F.S., Fe. S. and P.T developed the concept. F.C., J.Z. and J.B. designed and prepared samples. F.C. and J.Z. performed and analyzed measurements. P.T. supervised the project. All authors have written, read and approved the final manuscript.

