## Supplementary Information for "Super-Resolved FRET and Co-Tracking in pMINFLUX"

### Supplementary Section 1. pMINFLUX setup.

The pMINFLUX setup is described in the original pMINFLUX publication.[1] Depending on the excitation color, different optical elements such as filters, the vortex phase plate or polarization optics are used, however the beam path remains unchanged (see Supplementary Figure 1).

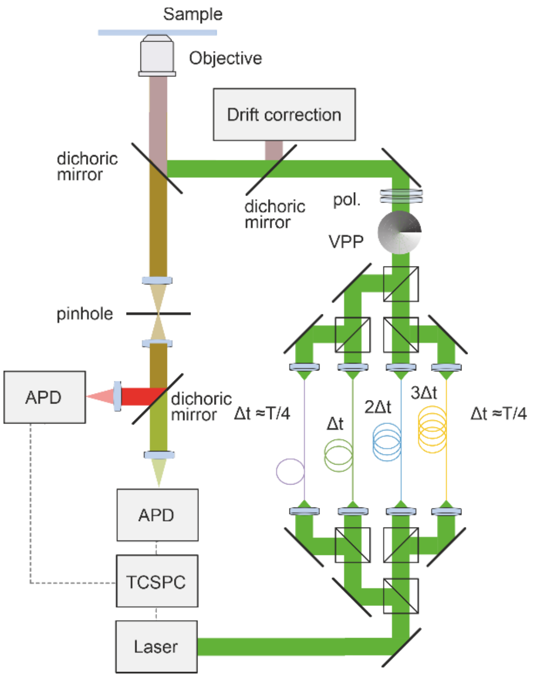


**Supplementary Figure 1: pMINFLUX setup.** A pulsed laser is split into four beams using beam splitters and coupled into optical fibers which delay the laser pulses as a function of the length of the fiber. The beams are recombined and doughnut-shaped beams are created with a vortex phase plate and polarization optics. The beams are focused onto the sample arranged in a triangular pattern with the fourth beam placed at the center of the triangle. For detection, APDs are used in combination with a TCSPC unit.

**Excitation.** A supercontinuum laser (SuperK Fianium FIU-15, NKT Photonics GmbH, Germany) is used at 19.5 MHz repetition rate as light source in combination with a tunable bandpass filter (SuperK VARIA, NKT Photonics GmbH, Germany) to select the desired wavelength range in the visible light spectrum. An additional clean-up filter (green: FLH532-10, Thorlabs GmbH, Germany, red: ZET 635/10, Chroma, USA) is used to further spectrally clean the excitation beam. Using a polarizing beam splitter cube (PBS251, Thorlabs GmbH, Germany), the light is split into two beams of orthogonal polarizations. Each of the beams is further split by a non-polarizing 50:50 beam splitter cube (BS013, Thorlabs GmbH, Germany). This beam splitting system generates two pairs of beams with each pair sharing the orthogonal linear polarization. The resulting four laser beams are coupled into polarization maintaining single-mode fibers (PM-S405-XP, Thorlabs GmbH, Germany) of lengths 2.0 m, 4.6 m, 7.1 m and 9.7 m such that the time delay between the beams after the fiber is ~12.5 ns (= T/4). The four beams are collimated after the fibers with an achromatic lens (AC254-035-A, Thorlabs GmbH, Germany) and recombined by using three 50:50 beam splitter cubes (BS013, Thorlabs GmbH, Germany). The overlay of the beams can be adjusted to obtain the required arrangement of laser foci in the object plane. The axes of linear polarization are matched by turning the fiber out-couplers (Thorlabs GmbH, Germany). Subsequently, the linearly polarized laser beams pass a combination of a quarter- and a half-wave plate (green: WPQ05M-532 and WPH532M532, Thorlabs GmbH, Germany; red: additional linear polarizer: LPVISC100-MP2, Thorlabs GmbH, Germany, RAC 5.2.10, B. Halle, Germany, WPQ05M-633, Thorlabs GmbH, Germany) to make them circularly polarized. A vortex phase plate (green: VPP, V-532-20-1, Vortex Photonics, Germany; red: VPP, V-633-20-1, Vortex Photonics, Germany) is then used to introduce the phase modulation necessary to generate the doughnut-shaped foci. The beams are guided into the back entrance of the microscope body (IX83, Olympus Deutschland GmbH, Germany), reflected on a dichroic mirror (ZT532/640rpc flat– STED, Chroma Technology Corp., USA) and focused with an objective (UPLSAPO100XO/1.4, Olympus Deutschland GmbH, Germany) onto the sample plane.

**Detection.** The fluorescence light is collected with the same objective and transmitted through the dichroic mirror, focused via an Olympus tube lens onto a pinhole (120 µm, Owis, Germany), collimated with an achromatic lens (AC254-150-A, Thorlabs GmbH, Germany) and spectrally split with a dichroic mirror (640 RDC dichroic mirror, Chroma, USA). The beams then are focused with a second achromatic lens (AC127-025-A, Thorlabs GmbH, Germany) to the chip of an avalanche photodiode (SPCM-AQRH-16-TR, Excelitas Technologies GmbH & Co. KG, Germany) after filtering the remaining scattered light from the laser with suitable interference optical filters (785 SP EdgeBasic, Semrock Inc., USA, green: 582/75 Brightline HC, Semrock Inc. USA, red: 700/75 ET Bandpass, Chroma, USA). The digital signal from the APD is sent to a TCSPC unit (HydraHarp 400, PicoQuant GmbH, Germany).

**Drift correction.** To measure and correct for sample drift during the measurement, the IR output of the variable bandpass filter is used. A beam of wavelength between 850 and 900 nm is selected with optical filters (875/50 bandpass, Edmund Optics GmbH), coupled into a single-mode fiber (780HP, Thorlabs GmbH, Germany), outcoupled and collimated. This beam is then split with a 50:50 beam splitter cube (BS014, Thorlabs GmbH, Germany) and combined again after inserting a lens system (ACN254-040-B, AC254-150-B, Thorlabs GmbH, Germany) into one of the two paths that focuses the beam to the back focal plane of the objective (dotted line) to create a widefield illumination at the sample plane. This beam is used for xy drift correction where the position of fiducial markers is localized during the measurement. The collimated IR beam is focused onto the sample plane at an oblique angle to achieve a z position-dependent spot at the detector and use this for z drift correction. Both IR beams are coupled to the main beam path via a dichroic mirror (ZT 785 SPXXR, Chroma Technology Corp., USA) and fed into the microscope to illuminate a region close, but not overlapping with the field of view used for MINFLUX. The reflected and backscattered light is split with an additional 50:50 beam splitter cube (BS014,

Thorlabs GmbH, Germany) from the excitation IR beam and detected on a single CMOS camera (Zelux, Thorlabs GmbH, Germany) at different positions of the chip.

**Setup control.** The piezo stage (P733.3CD, Physik Instrumente (PI) GmbH &Co. KG, Germany) translates the sample in all three dimensions with a resolution of 0.3 nm when running in closed loop mode. All components of the setup including the piezo stage are controlled digitally and integrated via a custom version of the PyFLUX project. Further details and source-code of this control software version are available at https://github.com/zaehringer-Jonas/pyflux

**Alignment.** For measurement the four vortex beams were aligned in a fixed triangular excitation beam pattern (EBP), with L ≈ 100-150 nm.

### Supplementary Section 2. FRET multilateration algorithm

#### Determination of FRET distances

The fluorescence lifetime of ATTO542 was determined via a fluorescence lifetime fit on the photons arriving at the green detection channel. The microtimes of these photons were extracted for each of the four excitation windows. They were then rebinned according to their maximal microtime bin in each excitation window and the resulting single histogram fitted with an IRF re-convoluted exponential fit. The fit model included an additional background component the weight of which was determined by a separate background measurement. By fitting each 'localization bin' in the time trace, the fluorescence lifetime of ATTO542 was extracted separately for each localization.

The FRET efficiency $E_{FRET}$ between two molecules was then calculated from the following equation:

$$E_{FRET} = 1 - \frac{\tau_{FRET}}{\tau_0}$$

where $\tau_{FRET}$ and $\tau_0$ are the fluorescence lifetimes of the FRET donor in presence and absence of the FRET acceptor, respectively. In the FRET pointer system, a fluorescence lifetime of $\tau_0$ = 3.3 ns was measured for ATTO542 in absence of the Cy5 acceptor.

As the FRET efficiency between two dye molecules depends on the distance $r$ between them with an inverse 6$^{th}$-power law, it can also be described as a function of this distance.

$$E_{FRET} = \frac{1}{1 + \left(\frac{r}{R_0}\right)^6}$$

where $R_0$, the so-called Förster distance, defines the inter-dye distance at which 50% energy transfer occurs. For the donor-acceptor pair ATTO542/ Cy5 in the FRET pointer system, we assumed a Förster distance of $R_0$ = 7.06 nm.

By combining both equations, the distance between ATTO542 and Cy5 was calculated for each MINFLUX localization using the corresponding measured fluorescence lifetime $\tau_{FRET}$.

$$r = R_0 \sqrt[6]{\frac{1}{\frac{\tau_0}{\tau_{FRET}} - 1}}$$

#### Multilateration of FRET acceptor position

The position of the red acceptor fluorophore in the FRET pointer measurement was multilaterated by combining each pMINFLUX localization with its respective FRET radius. For each localization of the ATTO542 donor fluorophore, a circle centered around the localization with a radius corresponding to the FRET radius of the localization was drawn. These circles were convoluted with a Gaussian distribution with a standard deviation corresponding to the uncertainties in position and radius of the circle. The uncertainty in position of the circle center was estimated by the precision of the ATTO542 localizations. The uncertainty in radius was calculated in an error propagation of the error of the fluorescence lifetime fit. The resulting density maps describe the probability of the FRET acceptor to be found at different positions, individually for each localization.

To multilaterally localize the FRET acceptor, multiple of these density maps were combined in a multiplicative fashion. The Gauss-convoluted FRET circles of all localizations were multiplied, resulting in the multiplicative density map shown in Fig, 1H.

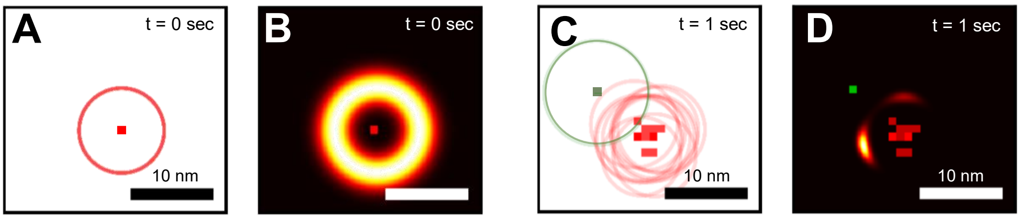


**Supplementary Figure 2: Multilateration of a FRET acceptor position by pMINFLUX.** By combining FRET donor (ATTO542) localizations (squares) with FRET distances (calculated through the corresponding fluorescence lifetimes of the FRET donor, circles), a probability density map for the location of the FRET acceptor (Cy5) was created. (A, C) FRET donor location and FRET distance combinations for (A) the first measurement point and (C) after 1 sec of measurement. (B,D) corresponding multiplicative probability density maps for the location of the FRET acceptor.

**Supplementary Section 3. pMINFLUX lifetime multiplexing algorithm and validation with single molecules**

**Model fit function**

The fluorescence decay of a single emitter $F(t)$ can be described by the probability density function (PDF) of an exponential distribution.

$$F(t) = \tau^{-1} \exp\left(-\frac{t}{\tau}\right) u(t)$$

where $t$ is the emission time of a photon with respect to the preceding laser pulse, $\tau$ denotes the fluorescence lifetime of the corresponding emitter and $u(t)$ is the unit step function of $t$.

To correct for the instrument response function (IRF) and temporal crosstalk due to the finite repetition period $T$ of pulsed lasers, the fluorescence decay is convolved with the IRF, and the fluorescence signal of the preceding excitation pulse added to the recorded fluorescence intensity.

$$I(t) = \sum_{i=0}^{1} F(t + iT, \tau) \star \mathrm{irf}$$

where $I(t)$ is the PDF of the measured fluorescence intensity, $\mathrm{irf}$ the normalized IRF and $\star$ the convolution operator.

In pMINFLUX, emitters are excited by four different, temporally delayed beam pulses per repetition period. The fit model thus is expanded to account for four beam pulses with the IRFs $\mathrm{irf_j}$.

$$I(t) = \sum_{j=0}^{3} a_j \sum_{i=0}^{1} F(t + iT, \tau) \star \mathrm{irf}_j$$

Here, $a_j$ describes the integrated fluorescence intensity upon excitation by beam *j*. To maintain normalization of the PDF, $a_j$ must be normalized to the accumulated integrated fluorescence intensity caused by all beams such that $\sum_j a_j \ = 1$.

To account for the fluorescence of multiple emitters, the fit model is expanded further to include the fluorescence decay of $K$ emitters of different lifetimes $\tau_k$.

$$\sum_{j=0}^{3} a_j \sum_{k=0}^{K} b_{jk} \sum_{i=0}^{1} F(t + iT, \tau_k) \star \mathrm{irf}_j$$

where $b_{jk}$ describes the integrated fluorescence intensity of emitter $k$ upon excitation with beam $j$. Similar to $a_j$, the ratios of $b_{jk}$ must be normalized to the accumulated integrated fluorescence intensity of all emitters excited by beam $k$ such that $\sum_k b_{jk} = 1$.

Contributions from background photons are included by adding a fraction $\gamma$ of background signal $\mathrm{bg}(t)$ added to the model fit function. The final model fit function thus is described by the following equation.

$$I(t) = (1-\gamma)\left[\sum_{j=0}^{3} a_j \sum_{k=0}^{K} b_{jk} \sum_{i=0}^{1} F(t+iT,\tau_k) \star \mathrm{irf}_j\right] + \gamma \cdot \mathrm{bg}(t)$$

Both $\mathrm{bg}(t)$ and the fraction $\gamma$ of $\mathrm{bg}(t)$ scatter as well as the fluorescence lifetimes $\tau_k$ were determined in separate background and calibration measurements. As a result, the fit function only depends on the parameter sets $a_j$ and $b_{jk}$. Due to the normalization constraints placed on the parameter sets, only a total number of $(4 \cdot K - 1)$ parameters must be fitted when describing the fluorescence of $K$ emitters in pMINFLUX measurements.

Parameter sets $a_j$ and $b_{jk}$. are retrieved separately for each localization by fitting the model fit function to the corresponding pMINFLUX TCSPC data using maximum likelihood estimation.

Multiplying the parameter sets $a_j$ and $b_{jk}$ then gives a set of four parameters for each emitter $k$ which describe the relative integrated fluorescence intensities of emitter $k$ upon excitation with laser beam pulses $j$. These integrated fluorescence intensities correspond to the photon counts calculated for emitter $k$ for the four beam pulses. They thus can be used as input parameters for the MINFLUX localisation algorithm, making the simultaneous localisation of multiple emitters possible.

### Supplementary Section 4. Performance of the fluorescence fitting approach for the localization of single emitters

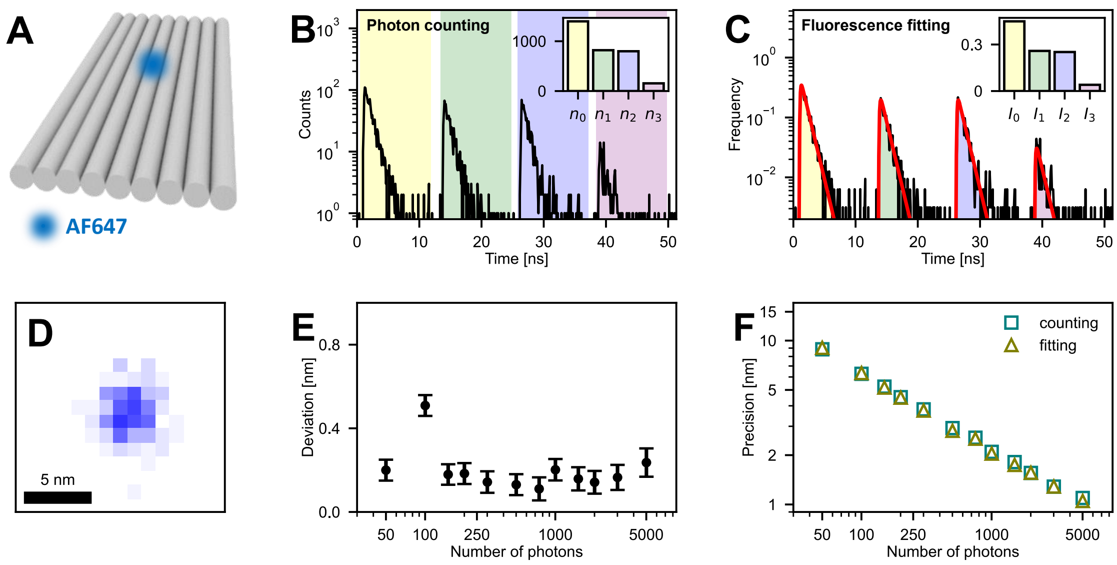


**Supplementary Figure 3: Comparison of the localization of a single emitter via fluorescence fitting and photon counting.** (A) Schematic of the static DNA origami carrying a single AF647 dye ($\tau$ = 1.1 ns) used in the pMINFLUX measurement to compare localizations obtained with the different approaches. (B, C) Fluorescence microtime decay of a pMINFLUX measurement of a single DNA origami as shown in (A). Colored areas mark (B) the detection microtime windows of the four different excitation beams and (C) the corresponding integrated fluorescence intensities extracted from the fitted fluorescence decay (red) used in the photon counting and the fluorescence fitting approach, respectively. The insets show (B) the photon numbers and (C) the relative fluorescence intensities upon excitation with the four different beams extracted from the pMINFLUX TCSPC data for both approaches. (D) 2D histogram of the pMINFLUX localizations of AF647 obtained by fluorescence fitting. (E) Deviation in position of localizations obtained by photon counting and fluorescence fitting as a function of the number of photons used for each localization. Deviations were calculated for the mean positions obtained from 2D Gaussian fits to 2D localization histograms as exemplarily shown in (D). (F) Localization precisions when using the fluorescence fitting and the photon counting approach as a function of the number of photons used for each localization.

### Supplementary Section 5. Excitation and emission spectra of AF647 and ATTO647N

The absorption and emission spectra of AF647 and ATTO647N attached to DNA are shown in Supplementary Fig. 4. The absorption spectra of both dyes are similar, whereas the emission spectrum of ATTO647N features a small red shift compared to AF647. Both the absorption spectrum of AF647 and the emission spectrum of ATTO647N as well as the absorption spectrum of ATTO647N and the emission spectrum of AF647 have a substantial overlap.

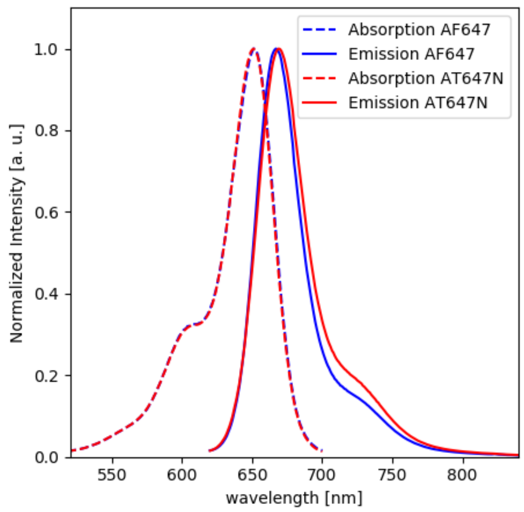


**Supplementary Figure 4: Normalized absorption and emission spectra of AF647 and ATTO647N covalently bound to DNA.**

**Supplementary Section 6. Performance of pMINFLUX lifetime multiplexing in presence of two, one and zero emitters.**

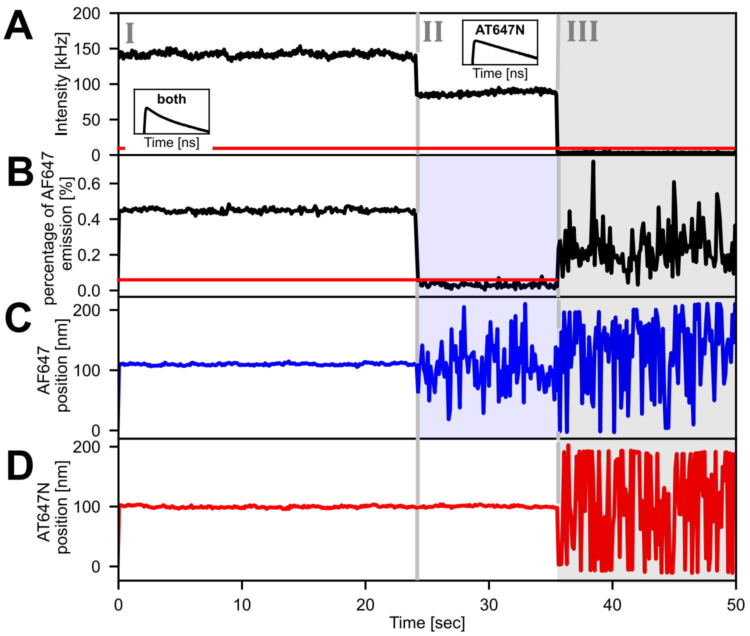


**Supplementary Figure 5: pMINFLUX lifetime multiplexing in presence of two, one and zero emitters.** Data analysis in this figure is based on the same measurement as in Fig. 2. (A) Fluorescence intensity transient recorded for a single DNA origami structure with AF647 and ATTO647N at a distance of 14.6 nm during a pMINFLUX measurement. Insets show the fluorescence lifetime decays before and after photobleaching of the first emitter, AF647 (first dashed gray line). Photobleaching of the second emitter, AT647N, and the subsequent drop of the fluorescence intensity to the background level is indicated by the second dashed gray line. The red line indicates a background threshold. Measurement points with a fluorescence intensity below this threshold are discarded (gray areas). (B) Percentage of AF647 emission of the total emission of both fluorophores as determined by the pMINFLUX lifetime multiplexing algorithm. In presence of both emitters, ~45% of the emitted photons are attributed to AF647 whereas the percentage drops close to zero after photobleaching of AF647. The red line indicates a percentage threshold for AF647. For measurement points with an AF647 emission percentage below this threshold, the localizations of AF647 are discarded as photobleaching of this emitter is assumed (blue areas). For AT647N, a threshold is defined analogously. (C) Position of AF647 as determined by pMINFLUX lifetime multiplexing. In presence of both emitters, AF647 is localized at a constant position. After photobleaching of AF647, its apparent localizations are distributed over the whole field of view (0-200 nm). However due to the priorly set thresholds, these false localizations are discarded. (D) Position of AT647N as determined by pMINFLUX lifetime multiplexing. Both in presence of both emitters and after photobleaching of AF647, the biexponential fitting approach of pMINFLUX lifetime multiplexing localizes AT647N at a constant position. Only after the second photobleaching step, the apparent localizations scatter. These false localizations however are discarded due to the priorly set background threshold.

**Supplementary Section 7. Bleach analysis in pMINFLUX**

The bleach analysis approach is based on similar concepts applied in wide-field super-resolution imaging.[2–4] Here, multiple emitters located within in a diffraction limited area are localized without photoswitching by imaging the same area multiple times. Ideally, during the imaging period the sequential photobleaching of all emitters occurs such that towards the end of the imaging period only one emitter is in its fluorescent state. This emitter can then be localized using standard procedures. For all images recorded prior to this, the fluorescence of the last emitter is then subtracted. This allows the localization of the emitter bleaching second-to-last. By again subtracting its fluorescence from all priorly recorded images, the next emitter can be localized. Repetition of this procedure eventually results in the full reconstruction of all emitter locations.

For its application in pMINFLUX, we adapted this concept to use the photon microtime information instead of recorded images. In the following, this adaption is described using the pMINFLUX measurement of AF647 and ATTO647N at a distance of 14.6 nm shown in Fig. 2 during which photobleaching of both emitters occurred (Supplementary Fig. 6A) as an example.

In a first step, we localized the emitter which photobleached last – in our case ATTO647N. For this, we applied the fluorescence fitting approach to the photon microtimes of photons detected after photobleaching of AF647 (time window II in Supplementary Fig. 6A) to extract the relative fluorescence intensities of ATTO647N upon excitation with the four different excitation beams (Supplementary Fig. 6B). We subsequently used these relative fluorescence intensities to determine the location of ATTO647N (red localization density map in Supplementary Fig. 6E) and calculated the number of photons emitted from ATTO647N by excitation with the different beams by multiplying the relative fluorescence intensities with the total number used for each localization.

We then applied the photon counting approach to the photon microtimes of photons detected while both AF647 and ATTO647N were in their fluorescent state (time window I in Supplementary Fig. 6A). The values extracted from this correspond to the number of photons emitted from both emitters upon excitation with the four different beams. To calculate the number of photons emitted from AF647 upon excitation with the different beams, we subtracted the number of photons emitted from ATTO647N as determined from time window II from the number of photons emitted from both emitters (see visualization in Supplementary Fig. 6C, Supplementary Fig. 6D). In a final step, the resulting photon numbers were used to determine the location of AF647 (blue localization density map in Supplementary Fig. 6E).

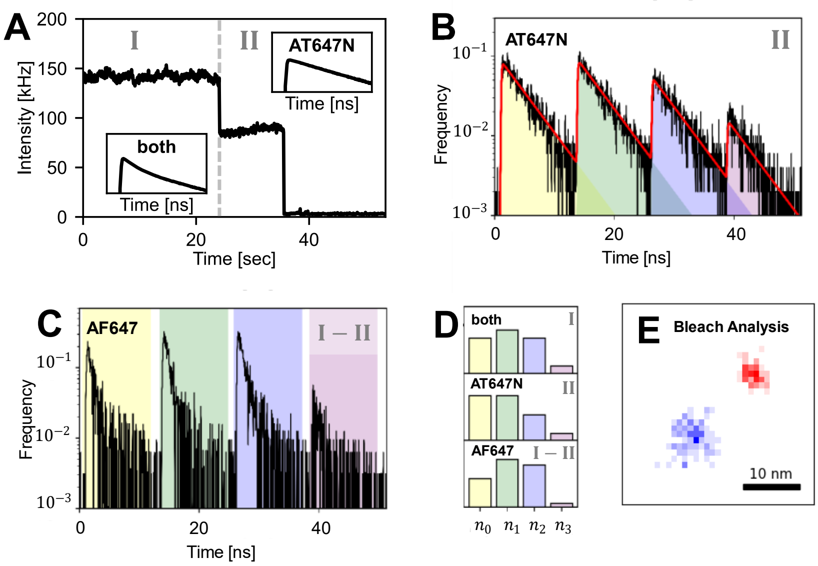


**Supplementary Figure 6: Bleach analysis for the sequential localization of multiple emitters in pMINFLUX without photoswitching.** The figure describes the bleach analysis used in Fig. 2 to validate the accuracy of the pMINFLUX multiplexing approach. Data analysis is based on the same measurement as in Fig. 2. (A) Fluorescence intensity transient recorded for a single DNA origami structure with AF647 and ATTO647N at a distance of 14.6 nm during a pMINFLUX measurement. Insets show the fluorescence lifetime decays before and after photobleaching of AF647 (dashed gray line). (B) Fluorescence microtime decay for ATTO647N in time window II in panel A. Colored areas mark the integrated fluorescence intensities for the different excitation beams extracted from the fluorescence fit (red line). The relative integrated fluorescence intensities then are used to localize ATTO647N. (C) Difference in the fluorescence microtime decays of time windows I and II, illustrating the sequential localization of AF647. Colored areas mark the detection microtime windows of the four different excitation beams in which the arriving photons are counted. (D) Retrieved normalized ratios of photon numbers/ fluorescence intensities of the four excitation beams for time window I (upper panel, fluorescence of both AF647 and ATTO647N), for time window II (middle panel, fluorescence of ATTO647N; used for the localization of ATTO647N) and for the difference between time windows I and II (lower panel, effective fluorescence of AF647; used for the localization of AF647). (E) 2D histogram of MINFLUX localizations obtained by bleach analysis. The localization of AF647 (blue) is less precise than the localization of ATTO647N (red) as it was not obtained directly.

### Supplementary Section 8. Effect of the fluorescent properties of the emitters on the performance of the pMINFLUX lifetime multiplexing approach

To estimate the effect of the fluorescent properties of the emitters on the overall performance of the pMINFLUX lifetime multiplexing approach, we performed different numerical simulations. Here, we compared the effects both different fluorescence lifetime contrasts as well as different brightness ratios have on the attainable localization precision.

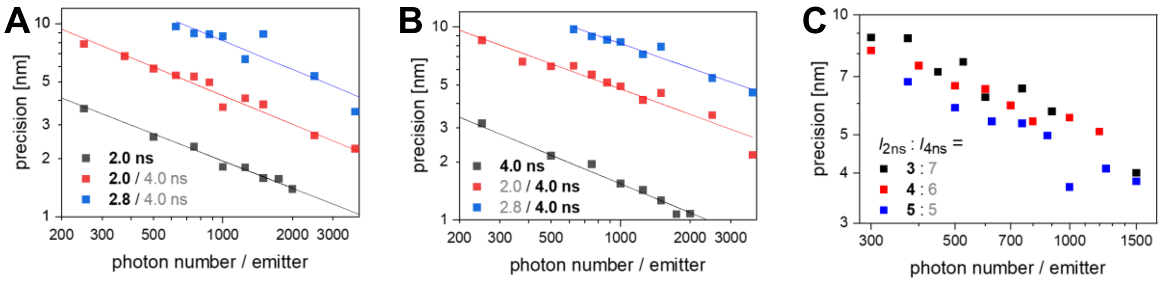


**Supplementary Figure 7: Effect of fluorescence lifetime contrasts and brightness ratios of two emitters on the performance of the pMINFLUX lifetimemultiplexing approach in numerical simulations.** (A,B) Localization precision as a function of the number of photons detected for the characterized emitter when simultaneously localizing two equally bright emitters of different fluorescence lifetimes by pMINFLUX lifetime multiplexing. The fluorescence lifetime of the characterized emitter is highlighted in black. For comparison, the localization precision when localizing a single emitter with the photon counting approach is given as reference. (C) localization precision of the emitter with a lifetime of $\tau_0$ = 2.0 ns as a function of the number of detected for the emitter when simultaneously localizing two emitters ($\tau_1$ = 4.0 ns) with different brightness ratios with pMINFLUX lifetime multiplexing. Numerical simulations were performed with a SBR of 10 and assuming a uniform background distribution.

### Supplementary Section 9. pMINFLUX lifetime multiplexing in the green spectral range

In the green spectral range, we identified ATTO542 and Alexa Fluor 555 (AF555) as a suitable emitter pair for pMINFLUX lifetime multiplexing. On reference DNA origami systems, ATTO542 exhibited a fluorescence lifetime of 3.3 ns whereas AF555 showed a biexponential decay profile with fluorescence lifetimes of 0.8 ns (85%) and 2.4 ns (15%), resulting in a high overall contrast in fluorescence lifetime of the emitter pair ideal for pMINFLUX lifetime multiplexing.

To account for the biexponential nature of the fluorescence of AF555, the model fit function of the multiplexing approach (see Supplementary Section 3) was extended to incorporate the biexponential decay of AF555 with fixed relative intensities.

We evaluated the performance of the pMINFLUX lifetime multiplexing approach when using AF555 and ATTO542 as an emitter pair by placing both emitters in a distance of ~18.7 nm from each other on a static DNA origami (Supplementary Fig. 8A) and performing a pMINFLUX measurement. The recorded fluorescence intensity/ fluorescence lifetime transient of the measurement (Supplementary Fig. 8B) featured time windows in which both fluorophores were in their fluorescent state (I), only AF555 was in its fluorescent state while ATTO542 was in a non-fluorescent state (II) and only ATTO542 was in its fluorescent state while AF555 was in a non-fluorescent state (III). This allowed localizing both AF555 and ATTO542 from time windows II and III using the standard photon counting approach (Supplementary Fig. 8C).

By applying the standard photon counting approach to photons arriving in time window I during which both fluorophores were in their fluorescent state, only the intensity-weighted average localization of both fluorophores was obtained (purple localizations in Supplementary Fig. 8C). In contrast, application of the pMINFLUX lifetime multiplexing approach to photons arriving in time window I revealed two separate locations for AF555 and ATTO542 (blue and red localizations in Supplementary Fig. 8D) which coincide with the positions for both fluorophores located via photon counting in Supplementary Fig. 8C, indicating a good accuracy of the pMINFLUX lifetime multiplexing approach when using AF555 and ATTO542 as an emitter pair.

When comparing the localization precisions achieved when simultaneously localizing both emitters using pMINFLUX lifetime multiplexing to those achieved when localizing single emitters using photon counting, localizations performed using pMINFLUX lifetime multiplexing are only less than two times less precise when using AF555 and ATTO542 as an emitter pair (Supplementary Fig. 8E,F).

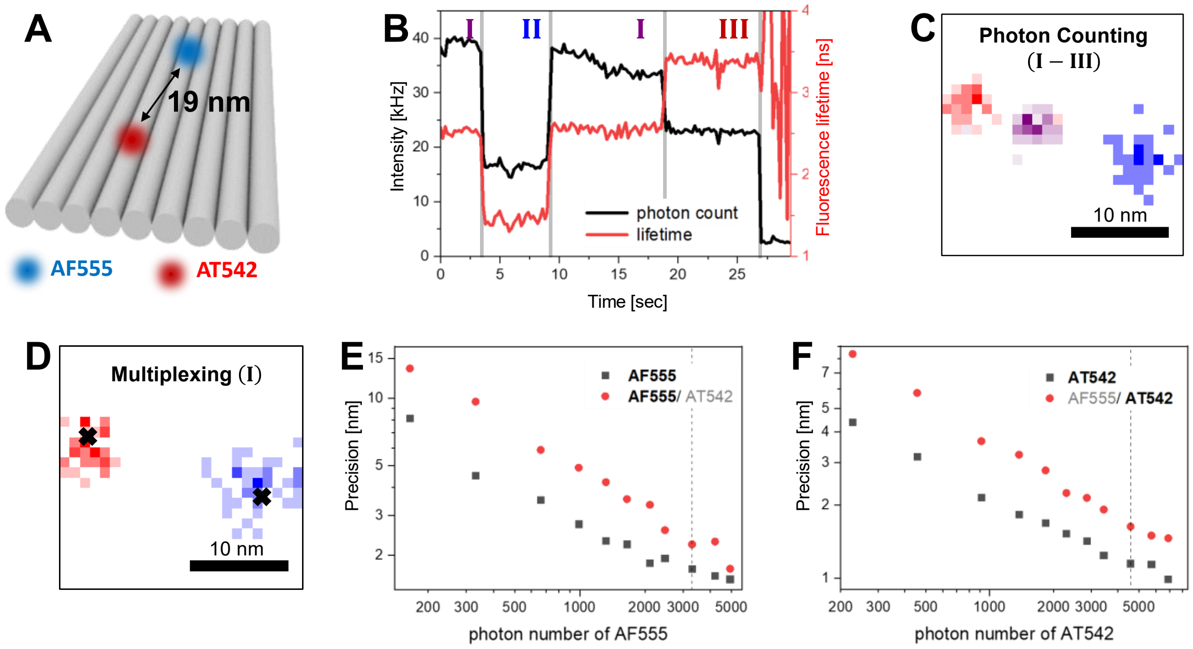


**Supplementary Figure 8: pMINFLUX lifetime multiplexing in the green spectral range using AF555 and ATTO542 as an emitter pair.** (A) Schematic of a static DNA origami on which both an AF555 dye and an ATTO542 dye are placed in a fixed distance of ~18.7 nm from each other. (B) Fluorescence intensity transient (black) recorded for a single DNA origami structure shown in panel A during a pMINFLUX measurement. The corresponding fluorescence lifetime transient as determined by a monoexponential fit model is shown in red. The dashed gray lines separate the transient into time windows during which both emitters (I) were in their fluorescent state and time windows during which only AF555 (II) or ATTO542 (III) were in their fluorescent state while the other emitter was in a non-fluorescent state. (C) 2D histogram of the pMINFLUX localizations in time windows I-III obtained by photon counting. Localizations of time window I corresponding to the combined fluorescence of both emitters are shown in purple, localizations of time window II corresponding to AF555 in blue and localizations of time window III corresponding to ATTO542 in red. (D) 2D histogram of the lifetime multiplexed pMINFLUX localizations in time window I. Localizations of AF555 are shown in blue, localizations of ATTO542 in red. The positions of the two emitters as determined by photon counting in panel C are indicated by black crosses. (E,F) Localization precision as a function of collected photons per emitter for both AF555 (E) and ATTO542 (F) localized simultaneously in time window I with pMINFLUX lifetime multiplexing (red). For comparison, the localization precision of the emitters localized during time windows II (E) and III (F) during which only the localized emitter was in its fluorescent state using the photon counting approach are shown in black. The dashed lines indicate the number of photons used per localization in the 2D histograms shown in panels C,D.

**Supplementary Section 10. Adjustment of the temporal resolution of the separate emitters in pMINFLUX lifetime multiplexing**

In pMINFLUX lifetime multiplexing, different temporal resolutions can be chosen for tracking the different emitters. If two processes with differing kinetics are studied, this allows separately optimizing the spatiotemporal resolution of both processes in a single, multiplexed measurement as demonstrated in Supplementary Fig. 9.

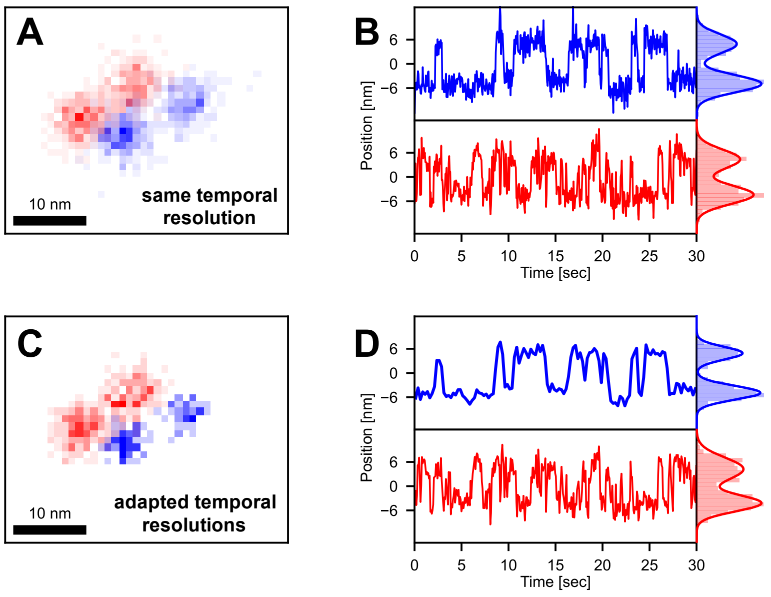


**Supplementary Figure 9: Dual-color molecular scale tracking of the double pointer system using pMINFLUX lifetime multiplexing with individually adaptable temporal resolutions.** (A,C) 2D histogram of the lifetime multiplexed pMINFLUX localizations of AF647 (blue) and ATTO647N (red), featuring each two distinct positions. (B,D) Localization trajectory of the AF647 (blue) and ATTO647N DNA pointer (red). In panels A and b, the temporal resolution is set to 60 ms for both colors. (C,D) The differing kinetics of the AF647 and the ATTO647N DNA pointer allows adjusting the temporal resolutions when tracking both pointer systems separately to 250 ms and 75 ms for the AF647 and the ATTO647N pointer, respectively, optimizing the spatiotemporal resolution separately for both tracked emitters.

**Supplementary Section 11. Monte-Carlo simulations on pMINFLUX lifetime multiplexing in the FRET range and Distance Calibration**

For pMINFLUX lifetime multiplexing, with AF647 and ATTO647N, two emitters with similar spectral properties are used. The resulting overlap in the absorption and emission spectra (see Supplementary Fig. 4) causes the occurrence of FRET, both from AF647 to ATTO647N and from ATTO647N to AF647 if the emitters are in close proximity to each other. To study the effect this has on the accuracy of the pMINFLUX lifetime multiplexing approach, we performed custom written kinetic Monte Carlo simulations simulating the fluorescence response of AF647 and ATTO647N placed at different distances from each other.

In these simulations, we assumed a gamma-shaped excitation pulse and an excitation probability $p_{ex}$ corresponding to the extinction coefficient of each emitter. Once excited, the emitter has multiple pathways. It can return to its non-fluorescent ground state either by fluorescence, by FRET or by a non-radiative pathway in probabilities described by the rates $k_f$, $k_{FRET}$ and $k_{nr}$, respectively.

The rates are coupled to photophysical properties such as the fluorescent lifetime $\tau$, the quantum yield of the fluorescent state $\phi$:

$$k_f = \tau_{noDNA}^{-1} \cdot \phi$$

where $\tau_{noDNA}$ is the fluorescence lifetime of the fluorophore without DNA modification. The modification with DNA only affects the non-radiative rate. The radiative rate remains the same thus can be calculated from manufacturer specifications. The non-radiative rate needs to be calculated from the experimental fluorescence lifetime with the corresponding buffer and DNA modification.

$$k_{nr} = \tau_{DNA}^{-1} - k_f$$

The FRET rate is strongly inter-dye distance r dependent as well as on the lifetime of the donor $\tau_D$ , as well as the Förster radius *R* characteristic for the dye pair:

$$k_{FRET} = \tau_D^{-1} \left(\frac{R_0}{r}\right)^6$$

As both dyes have similar spectral properties, both dyes can act as a donor hence, both FRET rates are needed.

Using the four rates for either dye (excitation, fluorescent emission, non-radiative relaxation and FRET), kinetic Monte-Carlo simulations[5] are performed for different distances. In the Monte-Carlo simulations we save where the photon is absorbed and where and when the photon is emitted. The kinetic Monte-Carlo framework is extended for pMINFLUX simulations. To this end, the excitation probability is modified according to the four vortex-beam excitation pattern and adding all four pMINFLUX excitations to one microtime window.

This framework is used to cross-check the experimental pMINFLUX lifetime multiplexing results with Monte-Carlo simulation for distances between 20 and 3 nm.

Rebinning the simulated photons according to their maximal microtime bin in each excitation window results in a single fluorescence decay (Supplementary Fig. 10A). The corresponding microtime decays show a distinct biexponential decay for large distances which becomes less distinct for smaller distances. In the simulations also the fraction of the photons absorbed and emitted by AF647 is determined (Supplementary Fig. 10B,C). As expected, the resulting fractions are strongly distance dependent and can be fitted by an adapted FRET relation:

$$E_{adapted} = E_{range} \cdot \left(1 + \left(\frac{d}{R_{50}}\right)^6\right)^{-1} + E_{low}$$

The fit results in $R_{50} = 7.8\,\mathrm{nm}$, $E_{range} = 19\%$ and $E_{low} = 48\%$ for the absorbed photons. This ratio is important to calculate the distance of the individual dyes to the center of mass of the MINFLUX measurement from the inter-dye distance. We then used photon packages of 2500 microtimes to calculate the phasor coordinates corresponding to the distances (Supplementary Fig. 10D,E). In simulations, the populations of AF647 and ATTO647N at distances of 10, 9, 8, 7, 6, 5 and 4 nm in the phasor plot can be easily differentiated. From of the phasor plot effective phasor coordinates can be calculated as the geometric mean of s and g.

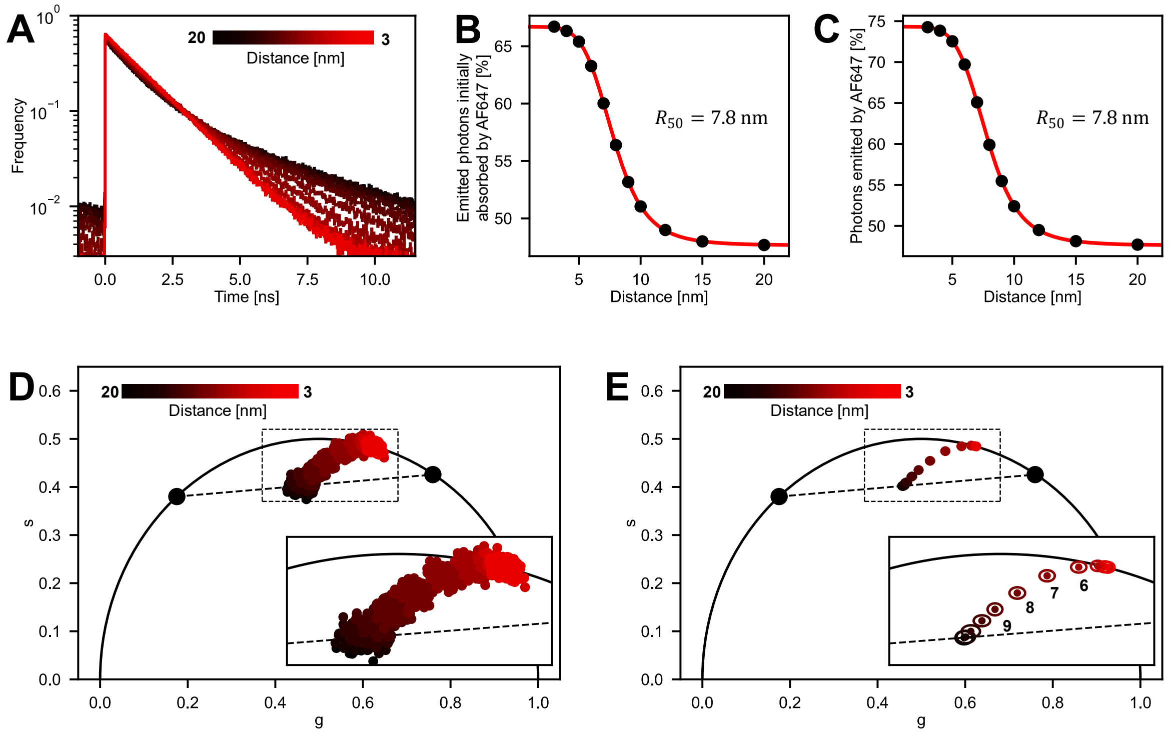


**Supplementary Figure 10: Phasor approach for distance determination in the FRET range – Monte Carlo simulations.** (A) Simulated fluorescence microtime decays of AF647 and ATTO647N placed at distances between 20 nm and 3 nm from each other in pMINFLUX experiments. The microtimes of the pMINFLUX simulations were extracted for each of the four excitation windows and then rebinned according to their maximal microtime bin in each excitation window, resulting in the shown single histogram (B,C) Fraction of photons absorbed (B) and emitted (C) by AF647 for different inter-fluorophore distances (black dots). Fitting with the adapted FRET equation revealed an apparent

Förster radius of $R_{50} = 7.8\,\mathrm{nm}$. (D, E) Simulated phasor plot of AF647 and ATTO647N placed at distances between 20 nm and 3 nm from each other in pMINFLUX experiments. The bold black dots correspond to the fluorescence of pure ATTO647N and AF647 (left and right dots), respectively. Datapoints on the dashed black line between them indicate inter-dye distances without interactions. The inset shows a zoom-in of the dashed box. Both a scatter plot of individual data points for all simulated distances (D) as well as a plot showing the mean phasor coordinates (dots) as well as the corresponding standard deviations (ellipses) for each simulated distance (E). The numbers in the inset of (E) correspond to the inter-dye distances in nanometer simulated for the respective data points. Simulations were carried out with a uniform background signal (overall SBR = 40) and using N = 2500 photons for calculating phasor data points in D,E.

The simulated data shows that the distance dependence of the phasor data follows the expected adapted FRET relation (Supplementary Fig. 11A). Using the same model for experimental data, also shows good agreement between fit and data. (Supplementary Fig. 11B). Thus, the fitted relation can be used as a calibration to calculate the distance out of the experimental phasor data used in Figure 4 (Supplementary Fig 11B black line).

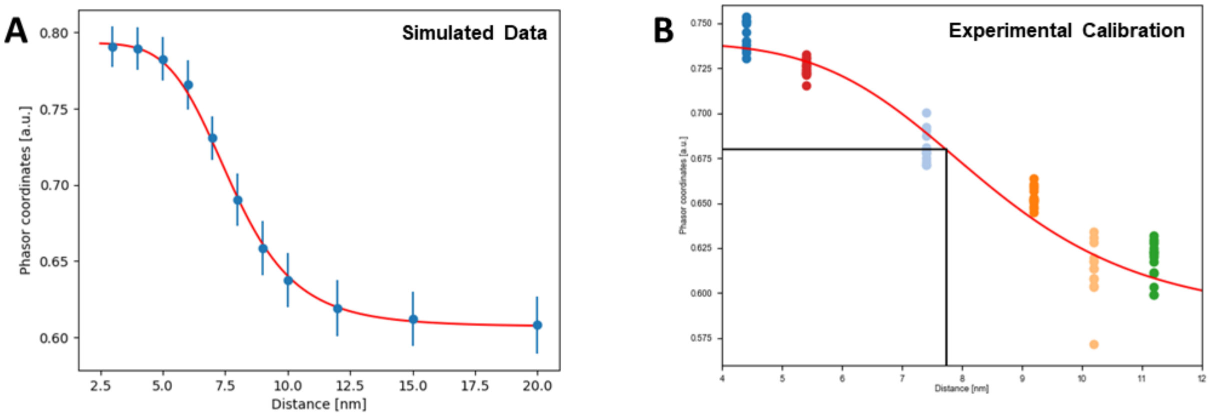


**Supplementary Figure 11: Calibration curve for distance determination via the phasor approach in the FRET range.** (A). The calibration curve of the simulated phasor coordinates and inter-dye distance. The red line represents the fit with the adapted FRET relation, used for calibration. (B). Color coded are the experimental phasor coordinates for samples of different inter-dye distances. The red line represents the fit with the adapted FRET relation, used for calibration. The black lines indicate the experimental measured phasor coordinate used in Figure 4 and its corresponding distance.

### Microtime-Gating in pMINFLUX

For localizations in which AF647 and ATTO647N were in distances smaller than ~10 nm, we exploited the microtime information offered in pMINFLUX to determine the vector along which both fluorophores were located. In the following, the concept is illustrated using Monte Carlo simulations.

An easy approach to partially separate the fluorescence of AF647 and ATTO647N is the application of two small microtime gates, directly after each pulsed excitation beam and at the end of the corresponding microtime window. As the fluorescence lifetimes of AF647 ($\tau$ = 1.1 ns) and ATTO647N ($\tau$ = 4.3 ns) differ, the early photons can be predominantly attributed to AF647 whereas the late photons mainly are emitted from ATTO647N.

We thus performed the standard pMINFLUX localization algorithm using only photons arriving in the respective early and late microtime gates, yielding two separate localizations. We then gradually increased the length of both microtime gates in steps of 250 ps to include larger fractions of photons emitted from both fluorophores (Supplementary Fig. 12A,B). With increasing microtime gate length, the resulting localizations are displaced further towards the center of mass of both fluorophores. Fitting a linear function to the localizations of the different microtime gates revealed correctly the vector defined by the actual positions of the two fluorophores (Supplementary Fig. 12C). This vector can be used in combination with the phasor distance. However, with smaller distance FRET gets stronger, and more photons get emitted by the fluorophore with lower fluorescence lifetime. Thus, in the MINFLUX one fluorophore is weighted more, moving the center of mass towards the fluorophore with lower fluorescence lifetime. This is directly proportional to the intensity ratio of Supplementary Figure 10B, which can be used to recover the dye-center of mass distance.

By combining all three parts: the vector connecting both dyes, the previously fitted phasor distance calibration (Supplementary Figure 11A) and the dye-center of mass distance, the individual fluorophores can be localized. In simulations the localized positions are in good agreement with the true positions, thus confirming the validity of the approach.

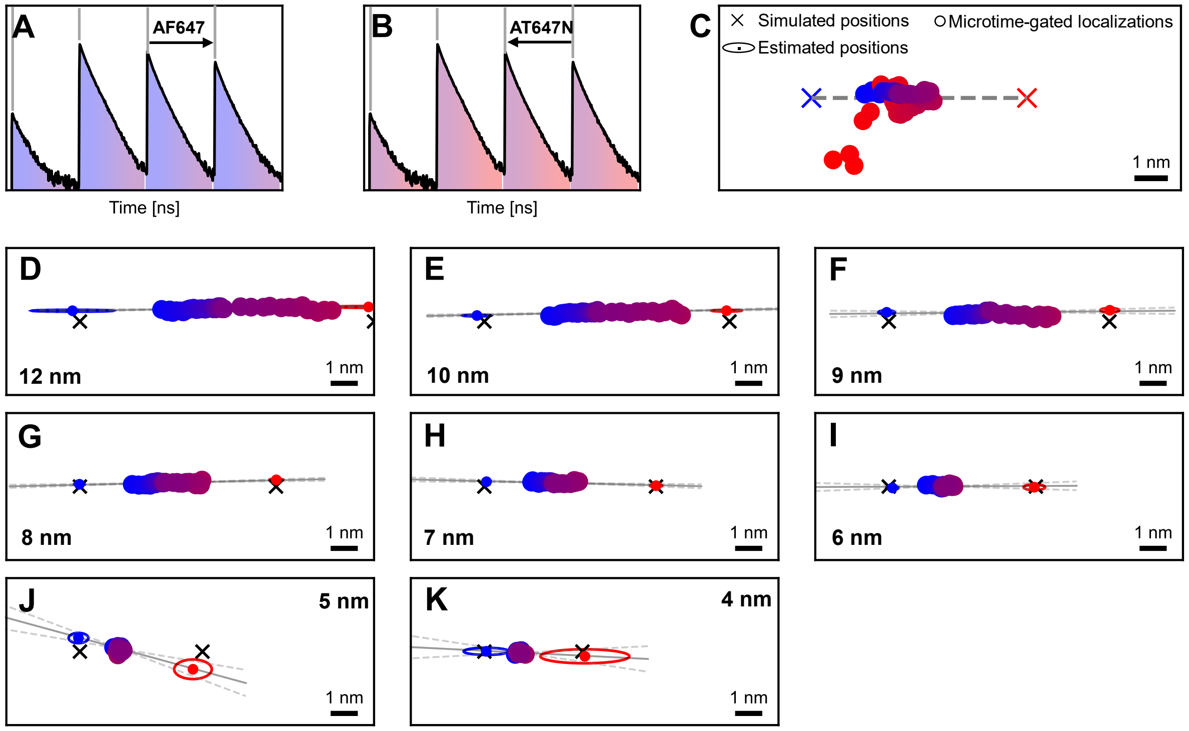

**Supplementary Figure 12: Microtime-Gating in the FRET range in pMINFLUX – Monte Carlo simulations.** (A, B) Simulated pMINFLUX microtime decays for AF647 and ATTO647N in a distance of 7 nm. For pMINFLUX localizations, only photons arriving in microtime gates right after each excitation pulse (A) or at the end of the corresponding microtime window (B) were considered. For determining the orientation of AF647 and ATTO647N to each other, the length of the microtime gates was gradually increased (color gradient to purple, black arrow). (C-K) Means of all localizations for the different gate lengths (circles) of the microtime decay shown in A,B (C) and AF647 and ATTO647N in distances between 12 nm and 4 nm. The color code corresponds to the different gate lengths illustrated in A,B. The large scattering in the mean localizations of late microtime gates in C (red circles) is caused by the increasingly shortened fluorescence lifetime of ATTO647N due to FRET and the resulting low number of photons/ high SBR in the late microtime gates. To circumvent this scattering, localizations performed with microtime gates filtering out more than 25% of the initial 8000 photons were discarded (D-K). The gray line indicates the fitted orientation vector. The dashed gray lines show the corresponding error margins. The blue and red dots and corresponding ellipses indicate the positions found for AF647 and AT647N with the phasor/ microtime-gating approach. Simulations were carried out with a uniform background signal (overall SBR = 40) and using N = 8000 photons for each localization. Initial microtime gate lengths were set to 0.25 ns and 3 ns for the early and the late photons, respectively. The gate lengths were increased in steps of 0.25 ns until they included the full microtime windows of the separate beam pulses. The SBR used for localizations was recalculated separately for each gate length to account for the different number of photons arriving in the different gates. For all distances, the microtimes of ~800 000 photons were simulated.

Due to the increasing indistinguishability of photons emitted from AF647 and ATTO647N due to FRET at small distances, the error of the determined vector direction; i.e. the slope of the linear fit, increases with decreasing distance (Supplementary Fig. 12D-K, Supplementary Tab. 1) The precision of the fit drops at distances of 5 nm and below, however still being accurate within 3 standard deviations.

**Supplementary Table 1: Slope of the Microtime-Gating Fit corresponding to the vector along which AF647 and ATTO647N are determined to be orientated.** The slope obtained from the ground truth of the simulations has a value of 0.

| Distance [nm] | Slope |
|---|---|
| 12 | 0.013 ± 0.005 |
| 10 | 0.019 ± 0.006 |
| 9 | 0.009 ± 0.019 |
| 8 | 0.022 ± 0.010 |
| 7 | -0.02 ± 0.014 |
| 6 | 0.005 ± 0.03 |
| 5 | -0.3 ± 0.10 |
| 4 | -0.05 ± 0.10 |

Experimental data was analyzed accordingly.

### Supplementary Section 12. Simulations on spectral multiplexing

An alternative approach to pMINFLUX multiplexing via the fluorescence lifetime utilizes the spectral information of the emitters. Here, emitters with similar absorption spectra but differences in their emission spectra are used. This allows exciting both emitters with the same excitation beam wavelength and still being able to partially separate their emission spectrally.

In spectral multiplexing, the fluorescence emission in MINFLUX experiments is spectrally split into two detection channels in each of which arriving photons are detected with an avalanche photon diode (APD0 and APD1, Supplementary Fig. 13A) using a dichroic mirror. The dichroic mirror optimally splits the fluorescence emission at a wavelength directly between the maxima in the emission spectra of the utilized emitters. This creates a contrast in the brightness with which both emitters are detected at the different APDs and allows their simultaneous localization.

In Supplementary Figure 13B the concept of spectral multiplexing is visualized using simulated pMINFLUX data of two emitters based on the fluorescent properties of ATTO542 and Cy3B which indicate that they could form a suitable emitter pair for spectral multiplexing in the green excitation range.

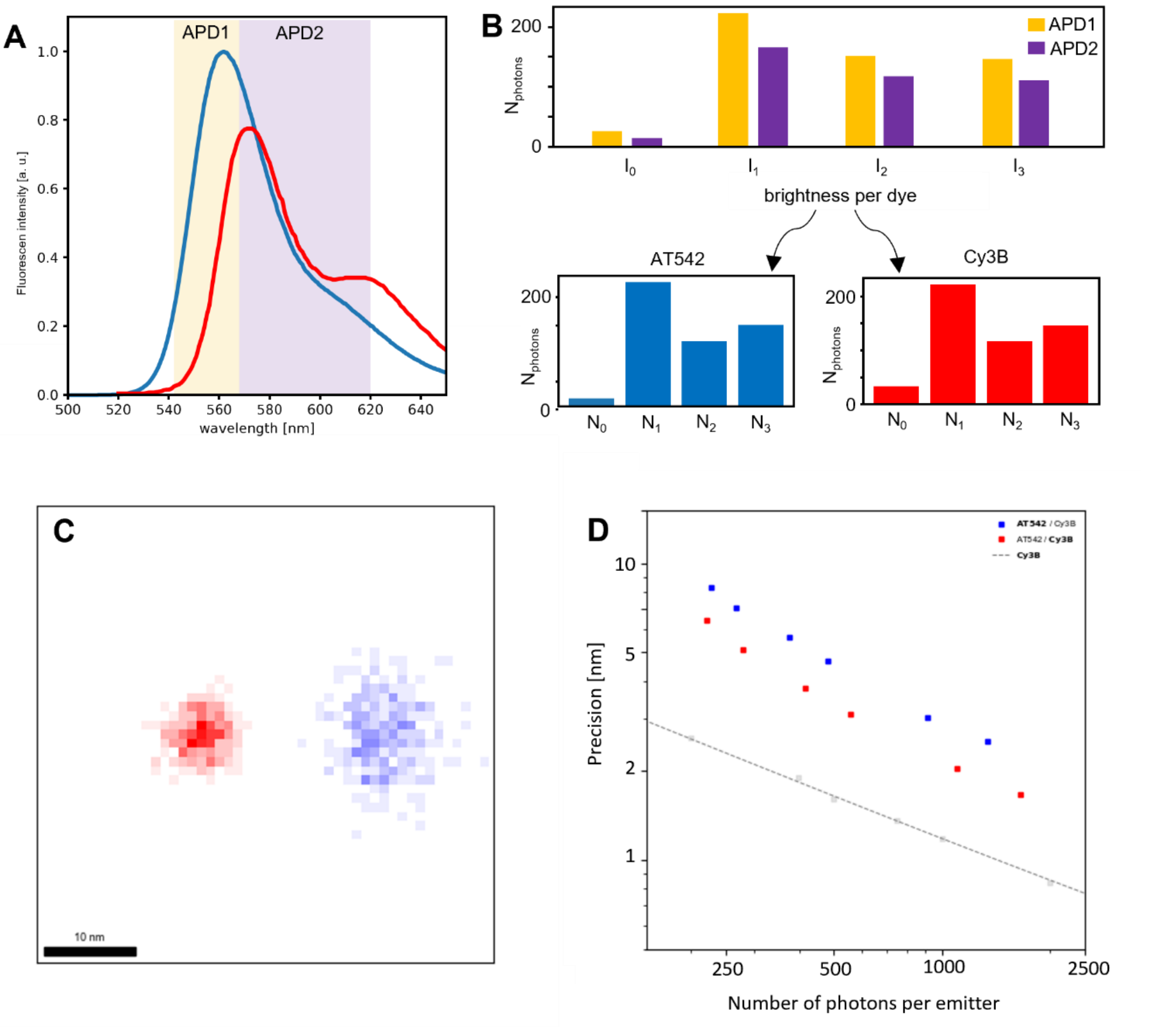


**Supplementary Figure 13: MINFLUX multiplexing via Spectral Splitting.** (A) Brightness normalized emission spectra of ATTO542 and Cy3b split with a dichroic mirror at 568 nm. The resulting spectral detection ranges of APD0 (yellow) and APD1 (purple) are marked. (B) Schematic of the spectral splitting workflow. For each localization, each excitation has detected photons on both APDs (yellow and purple). By knowing the brightness of each dye on each APD, the number of photons for each dye and each excitation can be extracted. (C) Simulated data of two dyes with spectral properties comparable to ATTO542 and Cy3B in a distance of 20 nm. Parameters for the simulations are 1000 localizations, with SBR = 50 and N = 2000 photons. Spectral splitting results in localizations of ATTO542 (blue) and Cy3B (blue) in a distance of 20 nm. (D) For simulations analogue to c, the precision of the spectral splitting localizations in dependence of the number of photons is comparable for both dyes (ATTO542: blue and Cy3B red). The precisions of a single dye (grey) at the same conditions outperforms the spectral splitting.

To extract the number of photons corresponding to each dye, the spectral splitting approach relies on knowing the fluorescent intensity fraction each dye has on each APD.

The total intensity on APD0 is the sum of the effective brightness of both dyes ($b_{dye,\#APD}$ with dye = 0,1 and # APD = 0,1), each multiplied with an intensity factor $\alpha_i$:

$$I_{APD\,0} = \alpha_0\, b_{0,0} + \alpha_1\, b_{1,0}$$

Similar for APD1:

$$I_{APD\,1} = \alpha_0\, b_{0,1} + \alpha_1\, b_{1,1}$$

The intensity factor is dependent on the position hence for each excitation beam different, but not on the APD channel.

Our effective measurement is the intensity ratio of both APDs:

$$r = \frac{I_{APD\,0}}{I_{APD\,1}} = \frac{\alpha_0\, b_{0,0} + \alpha_1\, b_{1,0}}{\alpha_0\, b_{0,1} + \alpha_1\, b_{1,1}}$$

$$\left(\alpha_0\, b_{0,1} + \alpha_1\, b_{1,1}\right) * r = \alpha_0\, b_{0,0} + \alpha_1\, b_{1,0}$$

$$\alpha_0\left(r\ b_{0,0} - b_{0,1}\right) = \alpha_1\left(b_{1,0} - r\ b_{1,1}\right)$$

$$\frac{\alpha_0}{\alpha_1} = \frac{b_{1,0} - r\ b_{1,1}}{r\ b_{0,0} - b_{0,1}}$$

By introducing the intensity fraction of each dye in the APDs $f_0 = \frac{b_{0,0}}{b_{0,0}+b_{0,1}} = \frac{b_{0,0}}{b_0}$ and $f_1 = \frac{b_{1,0}}{b_{1,0}+b_{1,1}} = \frac{b_{1,0}}{b_1}$ it follows:

$$\frac{\alpha_0}{\alpha_1} = \frac{f_1 b_1 - r * (1 - f_1) b_1}{r\, f_0 b_0 - (1 - f_0) * b_0} = \frac{f_1 - r + r f_1}{r\, f_0 - 1 + f_0} \frac{b_1}{b_0}$$

$$x = \frac{\alpha_0\, b_0}{\alpha_1\, b_1} = \frac{f_1 - r + r f_1}{r\, f_0 - 1 + f_0}$$

Here x is the absolute intensity ratio between both dyes.

$$x = \frac{N_0}{N_1}$$

It follows for the total number of photons for dye 0 ($N_0$):

$$N_0 = N - N_1 = N - \frac{N_0}{x} = \frac{x * N}{1 + x}$$

Using $N_0$ and $N_1$, the standard maximum likelihood approach of MINFLUX can be used. Hence only the expected fraction of a dye per APD needs to be known to calculate the number of photons corresponding to each dye out of the measured intensity ratio. This enables spectral multiplexing with a minimal number of parameters.

Next, we validated the spectral splitting approach in simulations. To this end two dyes spaced in a distance of 20.0 nm are simulated. Simulations parameter are a SBR of

50, 100 bins with each 2000 photons. The spectral splitting approach localized the dyes correctly in a distance of ~19.9 nm (Supplementary Fig. 13C). To characterize this method further simulations were performed, binning the localization with different number of photons. By measuring the precision for each of those simulation, the precision in dependence of the number of photons can be characterized analogue to Figure 3G (Supplementary Fig. 13D). The precision of the spectral splitting localizations is comparable for both dyes. The precision compared to simulations of a single dye with the same conditions shows a twofold reduction of the precision at the same number of photons. Compared to pMINFLUX multiplexing with a less than 1.5-fold reduction in precision at the same number of photons, spectral multiplexing is less photon efficient. However, spectral splitting is still considerably more photon efficient compared to widefield tracking methods. This makes spectral multiplexing attractive to implement in commercial continuous-wave MINFLUX setups.

### Supplementary Section 11. DNA staple strands used for folding the DNA origami structures

**Supplementary Table 2. Core staples from the 5‘ to the 3’ end for the L-shaped DNA origami structure used as a platform for the dynamic pointer systems.**

| Staple ID | Sequence (5’ to 3’) |
|---|---|
| 37[96] | CTCAAATGTTCAGAAATGGAAGTTTCACGCGCATTACTTCAACTGGCT |
| 23[120] | AAATTATTTGGAAACAGCCATTCGAAAATCGC |
| 26[171] | TAGTTGCCAGTTGCGGGAGGTTTTGAAGATCAATAA |
| 29[144] | AGCATGTACGAGAACAATCCGGTATTCTAAGAACGATTTTCCAGA |
| 35[112] | CATTATACGGTTTACCCATAACCCTCGAAATACAATGTTTAAACAGGG |
| 16[103] | CCCCCTGCGCCCGCTTTAGCTGTTTCCTGTGT |
| 18[167] | CAAACCCTTTAGTCTTACCAGCAGAAGATAA |
| 1[104] | GCAGCAAGCGGTCCACAAGTGTTTTGAGGCCA |
| 36[127] | ATTCATATCAGTGATTTGGCATCAGGACGTTGTAACATAAACCAGACG |
| 48[159] | AGGAGGTGGCGGATAAGTATTAAGAGGCTAAATCCTCTACAGGAG |
| 13[104] | CTAGCTGATAAATTAACAGTAGGG |
| 49[160] | CCAGAATGGAGCCGCCAATCAAGTTTGCC |
| 3[72] | CGTGCCTGTTCTTCGCATCCAGCGCCGGGTTA |
| 33[52] | TGTAGCTCAACATTTACCCTCGAAAGAC |
| 32[79] | ATCAAAAAGTCATAAAACGGAACAACATTATCAACTTTAGTAGAT |
| 12[119] | TATTTTTGAGAGATCTGCCATATTTCCTCTACTCAATTGA |
| 43[128] | CCGGAACCGCAAGAAAGCAATAGCTATCTTACTCACAATCCGATTGAG |
| 49[64] | CTGAGGCCAACGGCTACAGAGGTTTCCATT |
| 26[79] | ATAAAAATATCGCGTTCTCCTTTTGATAAGAGCTATAT |
| 15[88] | CAAAGGGCCTGTCGTGTGGCCCTGAGAGAGTT |
| 23[152] | ATTGCGTTTAACAACATTTCAATTACCTGAGCAAAAGGGAGAAACAGGTTTAAGAT GATGG |
| 32[111] | AGAAACAGCTTTAGAAGGAAGAAAAATCTACGATTTTAAGCATATAAC |
| 5[104] | GGGGTCATTGCAGGCGGGAATTGACTAAAATA |
| 34[172] | AGGAAACCGAGGACGTAGAAAAAGTACCG |
| 31[160] | GCCTAATTATCATATGATAAGAGATTTAGTTAATTTCAT |
| 42[63] | GACAGATGGACCTTCATCAAGAGCCCTGAC |
| 20[135] | TTATACTTAGCACTAAAAAGTTTGTGCCGCCA |
| 27[56] | GTTGTACCACCCTCATAAAGGCCGGAGACAG |
| 11[136] | GGCTTAGGTTGGGTTAAGCTAATGATTTTCGA |
| 2[87] | GAAATTGTTATCCGCTCACATTAAATTAATGA |
| 47[80] | CACAGACATTTCAGGGATCTCCAAAAAAAAGGTTCTTAAAGCCGCTTT |
| 9[72] | CCAGCCAGCTTTCCGGGTAATGGGGTAACAAC |
| 12[87] | CATTGCCTGAGAGTCTTTATGACCATAAATCATTTCATTT |
| 30[159] | CACTCATGAAACCACCTTAAATCAAGATTGAGCGTCTTTTTGTTT |
| 17[56] | ATGAGTGACCTGTGCAGTTTCTGCCAGCACG |
| 5[72] | CCAGCTTACGGCTGGAAACGTGCCCGTCTCGT |
| 16[135] | ACATTCTGAAGAGTCTCCGCCAGCAGCTCGAA |
| 51[104] | GGAGCCTTCACCCTCAGAGCCACC |
| 6[151] | AACGTTATTAATTTTACAACTAATCAGTTGGC |

| | |
|---|---|
| 8[151] | TGATTGCTTTGAATACAAACAGAATGTTTGGA |
| 18[135] | AAATCAACACGTGGCATCAGTATTCTCAATCC |
| 51[136] | TTGAGTAAGCCACCCTCAGAACCG |
| 19[120] | GTAAGAATAGTTGAAACTTTCGCAAACACCGC |
| 26[111] | CTTAATTGAGACCGGAAACAGGTCAGGATTAGAGGTGGCA |
| 15[56] | GTCCACTAAACGCGCGGACGGGCAACAGCTG |
| 32[143] | AACGTCAATAGACGGGGAATACCCAAAAGAACAAGACTCCGTTTTTAT |
| 22[135] | TTTTTTAATGCACGTACAAGTTACCCATTCAG |
| 20[167] | CAATTCATATAGATAATAAATCCTTTGCCCG |
| 23[56] | GTTTTCCCGTAGATGGCAGGAAGATCGCACT |
| 14[167] | GCCGATTAAGGAAGGGCGCGTAACCACCACA |
| 30[127] | TTTCATCGAATAATATCCAGCTACAATACTCCAGCAATTTCTTTACAG |
| 14[103] | CGGGAAACGAAAAACCTGATGGTGGTTCCGAA |
| 45[56] | GTCGAAATCCGCGACCTGCTCCACCAACTTTTAGCATTC |
| 10[87] | AAATCAGCTCATTTTTGTGAGCGAATAGGTCA |
| 21[152] | GCCGTCACAATATAAAAGAAACCACCAGAAGGAGCGGACTCGTATTACATTTGTC AAATAT |
| 43[96] | GGCACCAAAACCAAAAGTAAGAGCAACACTATAGCAACGTAAATCGCC |
| 24[103] | TAATAGTATTCTCCGTGCATTAAATTTTTGTT |
| 17[152] | AACCGTTTCACACGGGAAATACCTACATTTTGACGCTAAACTATCACTTCTTTAAC AGGAG |
| 46[172] | TTCTGAAACATGAAAGTGCCGGCCATTTG |
| 43[160] | CCTCAGAGCACAAGAAGAAAAGTAAGCAG |
| 49[96] | TGCGGGATAGCAGCGACGAGGCGCAGAGAAACGGCCGCGGTAACGATC |
| 50[79] | TACCGATAGTTGCGCTTTTTCA |
| 0[87] | ATCGGCAAAATCCCTTACGTGGACTCCAACGT |
| 12[151] | CTTCTGACCTAAATTTGCAGAGGCCAGAACGCAATTTACG |
| 9[136] | GCAGAGGCGAATTATTTTTCATTTGCTATTAA |
| 0[119] | AGGCGAAAATCCTGTTGTCTATCACCCCCGAT |
| 42[95] | AAGGGAACCGGATATTCACTCATCTTTGACCCGTAATGCCATCGGAAC |
| 4[87] | TGTTGCCCTGCGGCTGATCAGATGCAGTGTCA |
| 4[151] | AACAGAGGTGAGGCGGCAGACAATTAAAAGGG |
| 15[120] | TTAGAGCTATCCTGAGGCTGGTTTCAGGGCGC |
| 30[63] | TTAGTTTGCCTGTTTAGGTCATTTTTGCGGATAGGAAGCCGACTATTA |
| 42[127] | CCATTACCAAGGGCGACATCTTTTCATAGGCAGAAAGAATAGGTTGAG |
| 16[71] | CTGCGCGGCTAACTCACAATTCCACACAACATACGAGTACCGGGGCTCTGTGGG TGTTCAG |
| 45[128] | TCGATAGCAGCACCGTAAAATCACGTTTTGCT |
| 8[119] | GCTGCGCAACTGTTGGCAGACCTATTAGAAGG |
| 1[136] | GTACTATGGTTGCTTTTTTAGACACGCAAATT |
| 16[167] | TTCACCAGGTAGCAATGGCCTTGCTGGTAAT |
| 11[104] | GTATAAGCAAATATTTTAGATAAGTAACAACG |
| 9[104] | GGAAACCAGGCAAAGCGTACATAAGTGAGTGA |
| 19[56] | CAAATCGTCAGCGTGGTGCCATCCCACGCAA |
| 25[56] | TCATCAACAAGGCAAATATGTACCCCGGTTG |
| 24[135] | GCCTGTTTGCTTCTGTTACCTTTTAACGTTAA |
| 50[143] | TGTACTGGTAATAAGTTCAGTGCC |

| | |
|---|---|
| 18[103] | CACATCCTCAGCGGTGGTATGAGCCGGGTCAC |
| 47[112] | TAAAGTTTAGAACCGCTAATTGTATCGCGGGGTTTAAGTTTGGCCTTG |
| 3[104] | ACAGTTGAGGATCCCCAGATAGAACTGAAAGC |
| 21[120] | TCTTTAGGCTGAATAATGCTCATTAGTAACAT |
| 37[128] | AAGCGCATAAATGAAACAGATATAGAAGGCTTAGCAAGCCTTATTACG |
| 6[119] | GCAGTTGGGCGGTTGTCCAGTTATGGAAGGAG |
| 21[88] | CGCTGGCACCACGGGAGACGCAGAAACAGCGG |
| 27[88] | CTTTTGCGTTATTTCAATGATATTCAACCGTT |
| 39[51] | CAACTAATGCAGACAGAGGGGCAATACTG |
| 10[119] | TATTTTGTTAAAATTCGGGTATATATCAAAAC |
| 36[159] | AATAAGTTAGCAAAAACGCAATAATAACGAGAATTAAAAGCCCAA |
| 44[111] | GCACCCTCCGTCAGGTACGTTAGTAAATGAATAGTTAGCGTCAATCAT |
| 1[72] | ATTGCCCTTCACCGCCCCAGCTGCTTGCGTTG |
| 13[72] | TCAAATCACCATCAATACGCAAGG |
| 2[119] | TTCGTAATCATGGTCATCCATCAGTTATAAGT |
| 7[104] | ATCAAACTTAAATTTCTGGAAGGGCCATATCA |
| 6[87] | AAATCCCGTAAAAAAACGTTTTTTGGACTTGT |
| 48[127] | CCACCCTCTGTTAGGAAGGATCGTCTTTCCAGCAGACGATTATCAGCT |
| 51[168] | GCCCCCTGGTGTATCACCGTACTC |
| 35[80] | TGAATTACCAGTGAATGGAATTACGAGGCATATAGCGAGAGAATCCCC |
| 41[112] | AAAGACAAATTAGCAAGTCACCAATGAAACCA |
| 7[72] | CTCTCACGGAAAAAGAACGGATAAAAACGACG |
| 5[136] | CTGCAACAGTGCCACGTATCTGGTAGATTAGA |
| 22[167] | AATTACATAGATTTTCAATAACGGATTCGCC |
| 14[135] | ACGCCAGATGACGGGGCGCCGCTAGCCCCAGC |
| 3[136] | CAGGAAAAACGCTCATACCAGTAAATTTTTGA |
| 27[152] | GCCAGTACGTTATAAGGCGTTAAATAAGAATAAACACAAAT |
| 51[72] | CGTTGAAAATAGCAAGCCCAATA |
| 4[119] | GCCGGGCGCGGTTGCGCCGCTGACCCCTTGTG |
| 24[71] | TACAGGCATTAAATTAACCAATAGGAACGCCATCAAAGTCAATCAGAATTAGCCTAAATCG |
| 22[103] | AAACGGCGCAAGCTTTGAAGGGCGATCGGTGC |
| 19[88] | CCTGCAGCCATAACGGGGTGTCCAGCATCAGC |
| 20[71] | GAAACAACGCGGTCGCCGCACAGGCGGCCTTTAGTGACTTTCTCCACGTACAGACGCCAGG |
| 49[128] | ATATTCACCGCCAGCATTGACAGGCAAAATCA |
| 2[151] | ATCCAGAACAATATTAGTCCATCAGGAACGGT |
| 10[151] | CATAGGTCTGAGAGACAAATCGTCGAATTACC |
| 24[167] | ATAAACAATCCCTTAGTGAATTTATCAAAAT |
| 15[152] | GCGAGAAAAGGGATGACGAGCACGTATAACGTGCTTTTCACGCTGAAGAAAGC |
| 17[120] | CCGAGTAAGCCAACAGGGGTACCGCATTGCAA |
| 26[143] | TACCAGTAACGCTAACAGTTGCTATTTTGCACCCCATCCT |
| 50[111] | TGCTTTCGAGGTGAATCTCCAAAA |
| 19[152] | ATGGCTACAATCAACTGAGAGCCAGCAGCAAATGAAAAACGAACCTAATGCGCTTGGCAGA |
| 47[144] | CAGTACCATTAGTACCCAGTGCCCGTATAAATTGATGAATTAAAG |
| 37[160] | CCCTGAACAAATAAGAAACGCGAGGCGTT |

| | |
|---|---|
| 7[136] | TATCATTTTGCGGAACATCCTGATATAAAGAA |
| 48[63] | GGAACCCAAAACTACAAACAGTTTCAGCG |
| 27[120] | CCAACATGACGCTCAATGCCGGAGGAAATACC |
| 36[63] | GAGAAACATTTAATTTTACAGGTAGAAAG |
| 35[144] | CAGTATGTTTATTTTGCGAAGCCCTTTTTAATTGAGTTCTGAACA |
| 18[71] | AGAACGTTAACGGCGTAATGGGTAAAGGTTTCTTTGCGTCGGTGGTGCTGGTCTTGCCGTT |
| 41[144] | GGAGGGAAGAGCCAGCAATCAGTAGCGACAGACCAGAACCGCCTC |
| 51[51] | TGCGAATAATAATCGACAATGTTCGGTCG |
| 21[56] | CCGGCAAATCGGCGAAGTGGTGAAGGGATAG |
| 20[103] | TGGAGCCGGCCTCCGGGTACATCGACATAAAA |
| 31[128] | TCTTACCATAAAGCCATAATTTAGAATGGTTTAGGGTAGC |
| 8[87] | GGGCCTCTTCGCTATTACGTTGTACCTCACCG |
| 31[96] | AGCGAACCAGAAGCCTGGAGAATCACAAAGGCTATCAGGT |
| 36[95] | TGCTCATTCTTATGCGTTAATAAAACGAACTATATTCATTGGCTTTTG |
| 37[64] | CGGAATCTCAGGTCTGTTTTAAATATGCATGCGAACGAATCATTG |
| 23[88] | GCCAGTGCGATTGACCCACCGCTTCTGGTGCC |
| 25[152] | TTAATTTCATGTTCTATAACTATATGTAAATGCTGATGTCAATAGAATCCTTGACAAAATT |
| 48[95] | ACCCTCATGCCCTCATTTTCTGTATGGGATTTAGTTAAAGCAGCTTGA |
| 30[95] | AGTTGATTAGCTGAAAAGAGTACCTTTAATTGTTAATTCGGACCATAA |
| 17[88] | CGCTCACTATCAGACGGTCCGTGAGCCTCCTC |
| 29[112] | ACAAGAAATAGGAATCCCAATAGCAAGCAAATATAGCAGCATCCTGAA |
| 13[136] | GACCGTGTGATAAATACAAATTCT |
| 41[80] | ACAAGAACCGAACTGATGTTACTTAGCCGGAAAAGACAGCACTACGAA |
| 14[71] | ATCGGCCTTAAAGAATAAATCAAAAGAATAGCCCGAGACCAGTGAGGGAGAGGGGTGCCTA |
| 38[111] | ACGATAAACCTAAAACAAAGAATACACTAAAACATTACCCAACAAAGC |
| 42[159] | GGAATTAGGTAAATTTTCGGTCATAGCCCCACCGGAACCACCACC |
| 29[80] | GGGGCGCGCCCAATTCACTAAAGTACGGTGTCACGAGAATAGCTTCAA |
| 31[64] | TTCAAATTTTTAGAAAAAACAGGAGCAAACAAGAGAATCGATGAAGGGTGAGATATTTTA |
| 38[143] | TAATAAGAAGAGCCACCCTTATTAGCGTTTGCCATTCAACAATAGAAA |
| 25[120] | ATAACCTTATCAACAAAAATTGTATAACCTCC |
| 25[88] | CCGTCGGAGTAGCATTCAAAAACAGGAAGATT |
| 38[79] | CAAAAGAATAAAATACCCAGCGATTATACCAAGCGCGAA |
| 44[79] | GAGGGTAGTTGCAGGGTGCTAAACAACTTTCACGCCTGGAAAGAG |
| 22[71] | CGTTGGTAGTCACGACGCCAGCTGGCGAAAGGGGGATATCGGCCTGCGCATCGGCCAGCTT |
| 40[172] | TTTTCATCGGCATATTGACGGCACCACGG |
| 0[151] | CCCGCCGCGCTTAATGAAAGCCGGCGAACGTG |
| 44[143] | AGAGCCGCAAACAAATGAGACTCCTCAAGAGATTAGCGGGCAGTAGCA |
| 11[72] | ATAATCAGAAAAGCCCAACATCCACTGTAATA |
| 43[64] | AAACGGGGTTTTGCTACATAACGCCAAAAAAGGCTTGTAATCTTG |
| 3[25] | TTTTCGGGCCGTTTTCACGG |
| 41[36] | TTTTGGCGCATAGGCTGGCTAACGGTGTTAAATTGT |
| 45[173] | TTTCGACTTGATCGAGAGGGTTGATATAAGTATTTT |
| 45[31] | TTTTTATCATCGCCTGAACAGACCATTTT |

| | |
|---|---|
| 35[36] | TTTTATTGGGCTTGAGATGGCCAGAACGATT |
| 18[192] | TTTTACCTTGCTGAACCAGG |
| 19[179] | CTGATAGCCCTAAAACTTTT |
| 20[192] | TTTTTTCCTGATTATCACGT |
| 22[192] | TTTTAAACATCAAGAAAAAA |
| 4[178] | CCGAATCTAAAGCATCTTTT |
| 38[192] | TTTTGCTAATATCAGAGAGATAACCCCGCCACCGCG |
| 28[187] | TTTTCCCGACTTACAAAATAAACAGTTTT |
| 8[178] | TCGAAGATGATGAAACTTTT |
| 20[44] | CATGTTTACCAGTCCCTTTT |
| 15[176] | GAAAGGAGCGGGCGCTAGGTTTT |
| 11[25] | TTTTTTTTTTTTAAAACTAG |
| 43[36] | TTTTCTTTTTCACAACGGAGATTTGTTTT |
| 0[178] | CGGCCTCGTTAGAATCTTTT |
| 32[192] | TTTTCCATATTATTTATCCCAATCCAAAGTCAGAGA |
| 13[25] | TTTTGTGTAGGTAAAGATTC |
| 44[192] | TTTTCCCTCAGAGCCACCACCCTCAGAAAGCGCTTA |
| 8[198] | TTTTAACAGTACCTTTTACA |
| 0[198] | TTTTGCGCTGGCAAGTGTAG |
| 39[173] | ATACGCAAAGAAAATTATTCATTAAAGGTGAATTTT |
| 10[198] | TTTTGATTAAGACGCTGAGA |
| 14[192] | TTTTAGAGCGGGAGCTAGAT |
| 23[179] | CAGATGAATATACAGTTTTT |
| 14[44] | TTTGCGTATTGGGCGCTTTT |
| 40[187] | TTTTACTGTAGCCTCAGAACCGCCATTTT |
| 36[192] | TTTTCATATAAAAGAAAGCCGAACATTTT |
| 7[45] | AGATGAAGGGTAAAGTTTTT |
| 33[31] | TTTTATTGCTGAATATAATACATTTTTTT |
| 27[28] | TTTTGCCTCAGAGCATAAAGAAAATTAAGCAATAAATTTT |
| 2[178] | CTCCAATCGTCTGAAATTTT |
| 34[50] | ATTATAGCGTCGTAATAGTAAAATGTTTTTT |
| 18[44] | TCAGCAGCAACCGCAATTTT |
| 32[47] | TAGTCAGAAGCAAAGCGGATTTT |
| 10[178] | AGAGCAAATCCTGTCCAGATACCGACAAAAGGTAATTTT |
| 2[198] | TTTTTGCCTGAGTAGAAGAA |
| 37[36] | TTTTTAGACTGGCATCAGTTGAGATTTTTT |
| 17[31] | TTTTGTGTAAAGCCTGGCGG |
| 7[25] | TTTTGGAATTTGTGAGAGAT |
| 42[192] | TTTTTTATCACCGTCACAGCGTCAGTTTT |
| 49[36] | TTTTACGCATAATGAGAATAGAAAGTTTT |
| 29[36] | TTTTCGCAAATGGTCAATAAACCATTAGATGC |
| 12[200] | TTTTAGAACGCGAGAAAACTTT |
| 17[179] | TAGTAATAACATCACTTTTT |
| 21[179] | ATTTAGAAGTATTAGATTTT |
| 9[25] | TTTTTTGAGGGGACGACGAC |
| 11[45] | CATAATAATTCGCGTCTTTT |

| | |
|---|---|
| 13[45] | AAAACGGTAATCGTTTTTTT |
| 46[50] | GAGCCGATATAACAACAACCATCGCCCTTTTTTT |
| 16[44] | TGCGGCCAGAATGCGGTTTT |
| 5[25] | TTTTGAATGCCAACGGCAGC |
| 48[192] | TTTTTAGCCCGGAATAGCCTATTTCTTTT |
| 31[25] | TTTTTTGCATCAAAAGCCTGAGTAATTTT |
| 26[47] | AATGCAATAGATTAAGGGCTTAGAGCTTATTTT |
| 22[44] | CCGTGCATCTGCCAGTTTTT |
| 25[179] | ACATAGCGATAGCTTATTTT |
| 21[31] | TTTTTAAACGATGCTGATGG |
| 15[31] | TTTTTTGTTCCAGTTTGGAACAAGA |
| 50[192] | TTTTACCGTTCCAGTAAGCGTCATACATGGCTTCAGTTAAT |
| 5[45] | ACCTCGTCATAAACATTTTT |
| 12[178] | TTCCGGAATCATAATTTTTT |
| 47[36] | TTTTGTTTCGTCACCAGTACTGTACCGTAAT |
| 9[45] | AGTGTGCTGCAAGGCGTTTT |
| 4[198] | TTTTATCGCCATTAAAAATA |
| 46[187] | TTTTGGAACCTAAGTCTCTGAATTTTTTTTTT |
| 3[45] | TCACCGGAAGCATAAATTTT |
| 1[45] | TTCATAGGGTTGAGTGTTTT |
| 33[173] | TTAATTAAACCATACATACATAAAGGTGGCAATTTT |
| 26[192] | TTTTACTAGAAAAAGCCTGTT |
| 1[25] | TTTTCAGGGTGGTTTTTCTT |
| 23[31] | TTTTATTAAGTTGGGTACGC |
| 16[192] | TTTTTGGATTATTTACAGAA |
| 25[31] | TTTTTGGCCTTCCTGTATAA |
| 12[171] | ATATATATAAAGCGACGACATCGGCTGTCTTTCCTTATCATTTTT |
| 39[31] | TTTTTTAGGAATACCACAGTAGTAATTTT |
| 51[31] | TTTTGAACAACTAAAGGAACACTGATTTT |
| 24[195] | TTTTAGTAATTCAATCGCAAGACAATTTT |
| 30[192] | TTTTTCCAAGAACGGGTGCGAACCTTTTT |
| 19[31] | TTTTCCCTTACACTGGTTGC |
| 40[55] | ACAAAGTATGAGGAAGCTTTGAGGACTAAAGATTTT |
| 34[187] | TTTTAAGTTACCAGGGTAATTGAGCTTTT |
| 6[178] | ACAAATTATCATCATATTTT |
| 6[198] | TTTTCTTTACAAACAATTCG |

**Supplementary Table 3. Biotinylated staples for surface-immobilization of the L-shaped DNA origami structure.**

| Sequence (5‘ to 3‘) | Function |
|---|---|
| **biotin**-TACCAGTAACGCTAACAGTTGCTATTTTGCACCCCATCCT | immobilization |

| | |
|---|---|
| **biotin**-AGAGCCGCAAACAAATGAGACTCCTCAAGAGATTAGCGGG CAGTAGCA | immobilization |
| **biotin**-GAGGGTAGTTGCAGGGTGCTAAACAACTTTCACGCCTG GAAAGAG | immobilization |
| **biotin**-ATAAAAATATCGCGTTCTCCTTTTGATAAGAGCTATAT | immobilization |

**Supplementary Table 4. Staples for introducing the dynamic FRET pointer system with three protrusions in distances of 6 nm to each other to the L-shaped DNA origami structure.**

| Sequence (5‘ to 3‘) | Function |
|---|---|
| GGCACCAAAACCAAAAGTAAGAGCAACACTATAGCAACGTAAATCGCC TTTTTTTTTCGGGCATTTA-**ATTO542** | Pointer-ATTO542 at 3’ |
| AGAAACAGCTTTAGAAGGAAGAAAAATCTACGA TTTTA-Cy5 | Cy5 FRET acceptor |
| AACGAATCATTGTGAATTACCTTTTTTAAATGCC | Protrusion 1 |
| GGCACCAAAACCAAAAGTAAGAGCAACACTATAGCA ACTTTTTAAATGC | Protrusion 2 |
| AGCGTCAATCATAAGGGAACCGGTTTTAAATGCC | Protrusion 3 |
| GCACCCTCCGTCAGGTACGTTAGTAAATGAATAGTT | Exchange staple |
| TGCTCATTCAGTGAATGGAATTACGAGGCATATAGCG AGAGAATCCCC | Exchange staple |
| ATATTCACTCATCTTTGACCCGTAATGCCATCGGAAC | Exchange staple |
| ATATTCACCGCCAGCATCGATAGCAGCACCGTAAAAT CACGTTTTGCT | Exchange staple |
| CGGAATCTCAGGTCTGTTTTAAATATGCATGCG | Exchange staple |
| GTAAATCGCCAAAGACAAATTA | Exchange staple |
| GCAAGTCACCAATGAAACCATTGACAGGCAAAATCA | Exchange staple |
| ATGCGTTAATAAAACGAACTATATTCATTGGCTTTTG | Exchange staple |

**Supplementary Table 5. Staples for introducing the dynamic double pointer system with each two protrusions in a distance of ~12 nm to each other to the L-shaped DNA origami structure.**

| Sequence (5‘ to 3‘) | Function |
|---|---|
| GCTGCGCAACTGTTGGCAGACCTATTAGAAGGTGGAGCCGCCATTTTTT TTTCGGGCATTTA-**AF647** | AF647 pointer |

| | |
|---|---|
| ATCAAACTTAAATTTCTGGAAGTTTTTAAATGCC | AF647 protrusion |
| GGGCCTCTTCGCTATTACGTTGTACCTCACCGTTTTTAAATGCC | AF647 protrusion |
| GGCACCAAAACCAAAAGTAAGAGCAACACTATAGCAACGTAAATCGCC TTTTTTTTTCGGGCATTTA-**ATTO647N** | ATTO647N pointer |
| AAACGGGGTTTTGCTACATAACGCCAAAAAAGGCTTTTTTTTTAAATGC | ATTO647N protrusion |
| TGCCATTCAACAATAGAAAATTCATATGGTTTTTTTTTAAATGC | ATTO647N protrusion |
| TGCCCTGACGAGAAACATTTAATTTTACAGGTAGAAAG | Exchange staple |
| TAATAAGAAGAGCCACCCTTATTAGCGTT | Exchange staple |
| CATTATACCAGTGATTTGGCATCAGGACGTTGTAACATAAACCAGACG | Exchange staple |
| TTACCCATAACCCTCGAAATACAATGTTTAAACAGGG | Exchange staple |
| GACAGATGGACCTTCATCAAGAGTAATCTTG | Exchange staple |
| GGCCATATCAAAATTATTTGGAAACAGCCATTCGAAAATCGC | Exchange staple |
| GCCTCCGGGTACATCGACATAAAA | Exchange staple |
| CGGGAGACGCAGAAACAGCGG | Exchange staple |
| CCAGCTTACGGCTGGAAACGTGCCCGTCTCGTCGCTGGCA | Exchange staple |

**Supplementary Table 6. Core staples from the 5‘ to the 3’ end for the rectangular DNA origami structure used as a static platform for placing AF647 and ATTO647N in different distances to each other.**

| Staple ID | Sequence (5‘ to 3‘) |
|---|---|
| NRO-1-1 | CATAAATCTTTGAATACCAAGTGTTAGAAC |
| NRO-1-2 | GATGTGCTTCAGGAAGATCGCACAATGTGA |
| NRO-1-3 | GCAATTCACATATTCCTGATTATCAAAGTGTA |
| NRO-1-4 | GATTTAGTCAATAAAGCCTCAGAGAACCCTCA |
| NRO-1-5 | TCACCAGTACAAACTACAACGCCTAGTACCAG |
| NRO-1-6 | CCAATAGCTCATCGTAGGAATCATGGCATCAA |
| NRO-1-7 | GCTTTCCGATTACGCCAGCTGGCGGCTGTTTC |
| NRO-1-8 | AAAGGCCGGAGACAGCTAGCTGATAAATTAATTTTTGT |
| NRO-1-9 | AAATTAAGTTGACCATTAGATACTTTTGCG |
| NRO-1-10 | AAGCCTGGTACGAGCCGGAAGCATAGATGATG |
| NRO-1-11 | TCATTCAGATGCGATTTTAAGAACAGGCATAG |
| NRO-1-12 | GCCATCAAGCTCATTTTTTAACCACAAATCCA |
| NRO-1-13 | TATAACTAACAAAGAACGCGAGAACGCCAA |
| NRO-1-14 | TTGCTCCTTTCAAATATCGCGTTTGAGGGGGT |

| | |
|---|---|
| NRO-1-15 | GTATAGCAAACAGTTAATGCCCAATCCTCA |
| NRO-1-16 | AAAGTCACAAAATAAACAGCCAGCGTTTTA |
| NRO-1-17 | GGCCTTGAAGAGCCACCACCCTCAGAAACCAT |
| NRO-1-18 | TTAACGTCTAACATAAAAACAGGTAACGGA |
| NRO-1-19 | AGTATAAAGTTCAGCTAATGCAGATGTCTTTC |
| NRO-1-20 | TCAAATATAACCTCCGGCTTAGGTAACAATTT |
| NRO-1-21 | TTTCGGAAGTGCCGTCGAGAGGGTGAGTTTCG |
| NRO-1-22 | GAGGGTAGGATTCAAAAGGGTGAGACATCCAA |
| NRO-1-23 | TATTAAGAAGCGGGGTTTTGCTCGTAGCAT |
| NRO-1-24 | GCCCTTCAGAGTCCACTATTAAAGGGTGCCGT |
| NRO-1-25 | ATGCAGATACATAACGGGAATCGTCATAAATAAAGCAAAG |
| NRO-1-26 | AGCCAGCAATTGAGGAAGGTTATCATCATTTT |
| NRO-1-27 | TAAATGAATTTTCTGTATGGGATTAATTTCTT |
| NRO-1-28 | AAACAGCTTTTTGCGGGATCGTCAACACTAAA |
| NRO-1-29 | CGGATTCTGACGACAGTATCGGCCGCAAGGCGATTAAGTT |
| NRO-1-30 | GCGCAGACAAGAGGCAAAAGAATCCCTCAG |
| NRO-1-31 | AGAGAGAAAAAAATGAAAATAGCAAGCAAACT |
| NRO-1-32 | GACAAAAGGTAAAGTAATCGCCATATTTAACAAAACTTTT |
| NRO-1-33 | ACACTCATCCATGTTACTTAGCCGAAAGCTGC |
| NRO-1-34 | CTACCATAGTTTGAGTAACATTTAAAATAT |
| NRO-1-35 | TATATTTTGTCATTGCCTGAGAGTGGAAGATTGTATAAGC |
| NRO-1-36 | CGGATTGCAGAGCTTAATTGCTGAAACGAGTA |
| NRO-1-37 | TAAATCATATAACCTGTTTAGCTAACCTTTAA |
| NRO-1-38 | GTACCGCAATTCTAAGAACGCGAGTATTATTT |
| NRO-1-39 | TCTTCGCTGCACCGCTTCTGGTGCGGCCTTCC |
| NRO-1-40 | GCAAGGCCTCACCAGTAGCACCATGGGCTTGA |
| NRO-1-41 | ATTACCTTTGAATAAGGCTTGCCCAAATCCGC |
| NRO-1-42 | CTTATCATTCCCGACTTGCGGGAGCCTAATTT |
| NRO-1-43 | TTATACCACCAAATCAACGTAACGAACGAG |
| NRO-1-44 | GTAATAAGTTAGGCAGAGGCATTTATGATATT |
| NRO-1-45 | CAACCGTTTCAAATCACCATCAATTCGAGCCA |
| NRO-1-46 | GATGGTTTGAACGAGTAGTAAATTTACCATTA |
| NRO-1-47 | GCACAGACAATATTTTTGAATGGGGTCAGTA |
| NRO-1-48 | AGCAAGCGTAGGGTTGAGTGTTGTAGGGAGCC |
| NRO-1-49 | TCCACAGACAGCCCTCATAGTTAGCGTAACGA |
| NRO-1-50 | ATTATACTAAGAAACCACCAGAAGTCAACAGT |
| NRO-1-51 | TAAGAGCAAATGTTTAGACTGGATAGGAAGCC |
| NRO-1-52 | ATACATACCGAGGAAACGCAATAAGAAGCGCATTAGACGG |
| NRO-1-53 | CAACTGTTGCGCCATTCGCCATTCAAACATCA |
| NRO-1-54 | GATGGCTTATCAAAAAGATTAAGAGCGTCC |
| NRO-1-55 | TAGGTAAACTATTTTTGAGAGATCAAACGTTA |
| NRO-1-56 | AGGCAAAGGGAAGGGCGATCGGCAATTCCA |
| NRO-1-57 | ATTATCATTCAATATAATCCTGACAATTAC |
| NRO-1-58 | GAAATTATTGCCTTTAGCGTCAGACCGGAACC |
| NRO-1-59 | AATGGTCAACAGGCAAGGCAAAGAGTAATGTG |
| NRO-1-60 | ATACCCAACAGTATGTTAGCAAATTAGAGC |

| | |
|---|---|
| NRO-1-61 | ATAAGGGAACCGGATATTCATTACGTCAGGACGTTGGGAA |
| NRO-1-62 | CACCAGAAAGGTTGAGGCAGGTCATGAAAG |
| NRO-1-63 | ATCCCAATGAGAATTAACTGAACAGTTACCAG |
| NRO-1-64 | CATGTAATAGAATATAAAGTACCAAGCCGT |
| NRO-1-65 | CCAACAGGAGCGAACCAGACCGGAGCCTTTAC |
| NRO-1-66 | GCTATCAGAAATGCAATGCCTGAATTAGCA |
| NRO-1-67 | GACCTGCTCTTTGACCCCCAGCGAGGGAGTTA |
| NRO-1-68 | AGGAACCCATGTACCGTAACACTTGATATAA |
| NRO-1-69 | CAGCGAAACTTGCTTTCGAGGTGTTGCTAA |
| NRO-1-70 | ACAACTTTCAACAGTTTCAGCGGATGTATCGG |
| NRO-1-71 | CAGCAAAAGGAAACGTCACCAATGAGCCGC |
| NRO-1-72 | ACCTTTTTATTTTAGTTAATTTCATAGGGCTT |
| NRO-1-73 | CGATAGCATTGAGCCATTTGGGAACGTAGAAA |
| NRO-1-74 | GCCCGAGAGTCCACGCTGGTTTGCAGCTAACT |
| NRO-1-75 | ATTTTAAAATCAAAATTATTTGCACGGATTCG |
| NRO-1-76 | ACCTTGCTTGGTCAGTTGGCAAAGAGCGGA |
| NRO-1-77 | CTGAGCAAAAATTAATTACATTTTGGGTTA |
| NRO-1-78 | CCTGATTGCAATATATGTGAGTGATCAATAGT |
| NRO-1-79 | TCAATATCGAACCTCAAATATCAATTCCGAAA |
| NRO-1-80 | CTTTAGGGCCTGCAACAGTGCCAATACGTG |
| NRO-1-81 | AATAGTAAACACTATCATAACCCTCATTGTGA |
| NRO-1-82 | TCACCGACGCACCGTAATCAGTAGCAGAACCG |
| NRO-1-83 | GCCCGTATCCGGAATAGGTGTATCAGCCCAAT |
| NRO-1-84 | TGTAGCCATTAAAATTCGCATTAAATGCCGGA |
| NRO-1-85 | TCGGCAAATCCTGTTTGATGGTGGACCCTCAA |
| NRO-1-86 | TGACAACTCGCTGAGGCTTGCATTATACCA |
| NRO-1-87 | CCACCCTCTATTCACAAACAAATACCTGCCTA |
| NRO-1-88 | CCCGATTTAGAGCTTGACGGGGAAAAAGAATA |
| NRO-1-89 | AAGTAAGCAGACACCACGGAATAATATTGACG |
| NRO-1-90 | CACATTAAAATTGTTATCCGCTCATGCGGGCC |
| NRO-1-91 | TTAAAGCCAGAGCCGCCACCCTCGACAGAA |
| NRO-1-92 | ATATTCGGAACCATCGCCCACGCAGAGAAGGA |
| NRO-1-93 | TTCTACTACGCGAGCTGAAAAGGTTACCGCGC |
| NRO-1-94 | AACGTGGCGAGAAAGGAAGGGAAACCAGTAA |
| NRO-1-95 | GAATTTATTTAATGGTTTGAAATATTCTTACC |
| NRO-1-96 | AGCGCGATGATAAATTGTGTCGTGACGAGA |
| NRO-2-1 | AACGCAAAGATAGCCGAACAAACCCTGAAC |
| NRO-2-2 | GCCTCCCTCAGAATGGAAAGCGCAGTAACAGT |
| NRO-2-3 | AAAGCACTAAATCGGAACCCTAATCCAGTT |
| NRO-2-4 | GCCAGTTAGAGGGTAATTGAGCGCTTTAAGAA |
| NRO-2-5 | AAGGCCGCTGATACCGATAGTTGCGACGTTAG |
| NRO-2-6 | TTTTATTTAAGCAAATCAGATATTTTTTGT |
| NRO-2-7 | CTTTTGCAGATAAAAACCAAAATAAAGACTCC |
| NRO-2-8 | CCTAAATCAAAATCATAGGTCTAAACAGTA |
| NRO-2-9 | AGACGACAAAGAAGTTTTGCCATAATTCGAGCTTCAA |
| NRO-2-10 | AGAAAACAAAGAAGATGATGAAACAGGCTGCG |

| | |
|---|---|
| NRO-2-11 | CGCGCAGATTACCTTTTTTAATGGGAGAGACT |
| NRO-2-12 | CACAACAGGTGCCTAATGAGTGCCCAGCAG |
| NRO-2-13 | GCGGAACATCTGAATAATGGAAGGTACAAAAT |
| NRO-2-14 | TAAAAGGGACATTCTGGCCAACAAAGCATC |
| NRO-2-15 | AATTGAGAATTCTGTCCAGACGACTAAACCAA |
| NRO-2-16 | GCGAAAAATCCCTTATAAATCAAGCCGGCG |
| NRO-2-17 | AACACCAAATTTCAACTTTAATCGTTTACC |
| NRO-2-18 | TAAATCAAAATAATTCGCGTCTCGGAAACC |
| NRO-2-19 | GAAACGATAGAAGGCTTATCCGGTCTCATCGAGAACAAGC |
| NRO-2-20 | GCGAACCTCCAAGAACGGGTATGACAATAA |
| NRO-2-21 | TTAGGATTGGCTGAGACTCCTCAATAACCGAT |
| NRO-2-22 | ATCGCAAGTATGTAAATGCTGATGATAGGAAC |
| NRO-2-23 | GCGGATAACCTATTATTCTGAAACAGACGATT |
| NRO-2-24 | AAGGAAACATAAAGGTGGCAACATTATCACCG |
| NRO-2-25 | ACCCTTCTGACCTGAAAGCGTAAGACGCTGAG |
| NRO-2-26 | ATATTTTGGCTTTCATCAACATTATCCAGCCA |
| NRO-2-27 | TCAAGTTTCATTAAAGGTGAATATAAAAGA |
| NRO-2-28 | TCTAAAGTTTTGTCGTCTTTCCAGCCGACAA |
| NRO-2-29 | TTCCAGTCGTAATCATGGTCATAAAAGGGG |
| NRO-2-30 | AATACTGCCCAAAAGGAATTACGTGGCTCA |
| NRO-2-31 | TTTATCAGGACAGCATCGGAACGACACCAACCTAAAACGA |
| NRO-2-32 | TTGACAGGCCACCACCAGAGCCGCGATTTGTA |
| NRO-2-33 | CTGTGTGATTGCGTTGCGCTCACTAGAGTTGC |
| NRO-2-34 | GCGAGTAAAAATATTTAAATTGTTACAAAG |
| NRO-2-35 | TAGAGAGTTATTTTCATTTGGGGATAGTAGTAGCATTA |
| NRO-2-36 | CGAAAGACTTTGATAAGAGGTCATATTTCGCA |
| NRO-2-37 | TCATCGCCAACAAAGTACAACGGACGCCAGCA |
| NRO-2-38 | TTAACACCAGCACTAACAACTAATCGTTATTA |
| NRO-2-39 | TTATTACGAAGAACTGGCATGATTGCGAGAGG |
| NRO-2-40 | ACAACATGCCAACGCTCAACAGTCTTCTGA |
| NRO-2-41 | CATTTGAAGGCGAATTATTCATTTTTGTTTGG |
| NRO-2-42 | TGAAAGGAGCAAATGAAAAATCTAGAGATAGA |
| NRO-2-43 | TGGAACAACCGCCTGGCCCTGAGGCCCGCT |
| NRO-2-44 | TACCGAGCTCGAATTCGGGAAACCTGTCGTGCAGCTGATT |
| NRO-2-45 | GTTTATTTTGTCACAATCTTACCGAAGCCCTTTAATATCA |
| NRO-2-46 | ACAAACGGAAAAGCCCCAAAAACACTGGAGCA |
| NRO-2-47 | GTTTATCAATATGCGTTATACAAACCGACCGTGTGATAAA |
| NRO-2-48 | ACGGCTACAAAAGGAGCCTTTAATGTGAGAAT |
| NRO-2-49 | GACCAACTAATGCCACTACGAAGGGGGTAGCA |
| NRO-2-50 | CTCCAACGCAGTGAGACGGGCAACCAGCTGCA |
| NRO-2-51 | ACCGATTGTCGGCATTTTCGGTCATAATCA |
| NRO-2-52 | CAGAAGATTAGATAATACATTTGTCGACAA |
| NRO-2-53 | TGCATCTTTCCCAGTCACGACGGCCTGCAG |
| NRO-2-54 | TTAGTATCACAATAGATAAGTCCACGAGCA |
| NRO-2-55 | GTTTTAACTTAGTACCGCCACCCAGAGCCA |
| NRO-2-56 | TTAATGAACTAGAGGATCCCCGGGGGGTAACG |

| | |
|---|---|
| NRO-2-57 | CTTTTACAAAATCGTCGCTATTAGCGATAG |
| NRO-2-58 | ATCCCCCTATACCACATTCAACTAGAAAAATC |
| NRO-2-59 | AGAAAGGAACAACTAAAGGAATTCAAAAAAA |
| NRO-2-60 | AGCCACCACTGTAGCGCGTTTTCAAGGGAGGGAAGGTAAA |
| NRO-2-61 | AACAAGAGGGATAAAAATTTTTAGCATAAAGC |
| NRO-2-62 | GCCGTCAAAAAACAGAGGTGAGGCCTATTAGT |
| NRO-2-63 | TGTAGAAATCAAGATTAGTTGCTCTTACCA |
| NRO-2-64 | GAGAGATAGAGCGTCTTTCCAGAGGTTTTGAA |
| NRO-2-65 | CCACCCTCATTTTCAGGGATAGCAACCGTACT |
| NRO-2-66 | CTTTAATGCGCGAACTGATAGCCCCACCAG |
| NRO-2-67 | CCAGGGTTGCCAGTTTGAGGGGACCCGTGGGA |
| NRO-2-68 | CAAATCAAGTTTTTTGGGGTCGAAACGTGGA |
| NRO-2-69 | ACGCTAACACCCACAAGAATTGAAAATAGC |
| NRO-2-70 | TACGTTAAAGTAATCTTGACAAGAACCGAACT |
| NRO-2-71 | TAATCAGCGGATTGACCGTAATCGTAACCG |
| NRO-2-72 | TTTTCACTCAAAGGGCGAAAAACCATCACC |
| NRO-2-73 | GCCTTAAACCAATCAATAATCGGCACGCGCCT |
| NRO-2-74 | AATAGCTATCAATAGAAAATTCAACATTCA |
| NRO-2-75 | CATCAAGTAAAACGAACTAACGAGTTGAGA |
| NRO-2-76 | CAGGAGGTGGGGTCAGTGCCTTGAGTCTCTGAATTTACCG |
| NRO-2-77 | AAATCACCTTCCAGTAAGCGTCAGTAATAA |
| NRO-2-78 | CTCGTATTAGAAATTGCGTAGATACAGTAC |
| NRO-2-79 | TTTACCCCAACATGTTTTAAATTTCCATAT |
| NRO-2-80 | GTCGACTTCGGCCAACGCGCGGGGTTTTTC |
| NRO-2-81 | CGTAAAACAGAAATAAAAATCCTTTGCCCGAAAGATTAGA |
| NRO-2-82 | AGGCTCCAGAGGCTTTGAGGACACGGGTAA |
| NRO-2-83 | GAGAAGAGATAACCTTGCTTCTGTTCGGGAGAAACAATAA |
| NRO-2-84 | TTTAGGACAAATGCTTTAAACAATCAGGTC |
| NRO-2-85 | AATACGTTTGAAAGAGGACAGACTGACCTT |
| NRO-2-86 | CTTAGATTTAAGGCGTTAAATAAAGCCTGT |
| NRO-2-87 | TAAATCGGGATTCCCAATTCTGCGATATAATG |
| NRO-2-88 | AACAGTTTTGTACCAAAAACATTTTATTTC |
| NRO-2-89 | CTGTAGCTTGACTATTATAGTCAGTTCATTGA |
| NRO-2-90 | AACGCAAAATCGATGAACGGTACCGGTTGA |

**Supplementary Table 7. Staples from the 5‘ to the 3’ end for introducing AF647 and ATTO647N statically placed at different distances to each other on the rectangular DNA origami. The distances labelled 4, 5, 7, 9, 10, 11 and 14 correspond to inter-fluorophore distances of 4.4 nm, 5.4 nm, 7.4 nm, 9.2 nm, 10.2 nm, 11.2 nm and 14.6 nm, respectively.**

| Sequence (5‘ to 3‘) | Function |
|---|---|
| TTAAAGCCAGAGCCGCCACCCTCGACAGAAT-**AF647** | AF647 for distances 4, 5, 7 |

| | |
|---|---|
| **AF647**-TCCTTTAGCGTCAGACCGGAACC | AF647 for distances 9, 10, 11, 14 |
| TCAGAAACCATCGATAGCAGCACCGTAATC-**ATTO647N** | ATTO647N for distance 4 |
| TCAGAAACCATCGATAGCAGCACCGTA-**ATTO647N** | ATTO647N for distances 5, 9 |
| CAGCAAAAGGAAACGTCACCAATGAAACCATCGATAGCAGC-**ATTO647N** | ATTO647N for distance 7,11 |
| TCAGAAACCATCGATAGCAGCACC-**ATTO647N** | ATTO647N for distance 10 |
| GGCCTTGAAGAGCCACCACCCTCAGAAACCAT-**ATTO647N** | ATTO647N for distance 14 |
| GTGCCGTCGAGAGGGTGAGTTTCG | Exchange staple for distances 4,5,7, 9, 10,11 |
| AACAAATACCTGCCTATTTCGGAA | Exchange staple for distances 4,5,7, 9, 10,11 |
| AAGGAAACATAAAGGTGGCAACATTATCA | Exchange staple for distances 4,5,7, 9, 10,11 |
| GAAATTATTCATTAAAGGTGAATATAAAAGA | Exchange staple for distances 9, 10, 11, 14 |
| CCTCGACAGAATCAAGTTTG | Exchange staple for distances 9, 10, 11, 14 |
| TTAAAGCCAGAGCCGCCAC | Exchange staple for distances 9, 10, 11, 14 |
| AGTAGCAGAACCGCCACCCTCTATTCACA | Exchange staple for distances 4, 7, 11 |
| CCGTCACCGACTTGAGCCATTTGGGAACGTAGAAA | Exchange staple for distances 4, 5, 9, 10 |
| GGCCTTGAAGAGCCACCACCC | Exchange staple for distances 4, 5, 9, 10 |
| ATCAGTAGCAGAACCGCCACCCTCTATTCACA | Exchange staple for distances 5, 9 |
| GTAATCAGTAGCAGAACCGCCACCCTCTATTCACA | Exchange staple for distance 10 |
| ACCGTAATCCCGTCACCGACTTGAGCCATTTGGGAACGTAGAAA | Exchange staple for distances 7, 11 |
| GGCCTTGAAGAGCCACCACCCTCAGAGCCGC | Exchange staple for distances 7, 11 |

| | |
|---|---|
| **biotin**-CGGATTCTGACGACAGTATCGGCCGCAAGGCGATTAAGTT | immobilization |
| **biotin**-ATAAGGGAACCGGATATTCATTACGTCAGGACGTTGGGAA | immobilization |
| **biotin**-GAGAAGAGATAACCTTGCTTCTGTTCGGGAGAAACAATAA | immobilization |
| **biotin**-TAGAGAGTTATTTTCATTTGGGGATAGTAGTAGCATTA | immobilization |
| **biotin**-AGCCACCACTGTAGCGCGTTTTCAAGGGAGGGAAGGTAAA | immobilization |
| **biotin**-GAAACGATAGAAGGCTTATCCGGTCTCATCGAGAACAAGC | immobilization |
